# Structural and phenotypic plasticity of the RBD loop2 region is a key determinant for HKU5r-CoVs’ emergence in mink

**DOI:** 10.1101/2025.10.22.684003

**Authors:** Jarel Elgin Tolentino, Victoria A. Jefferson, Nicholas J. Catanzaro, Ralph S. Baric, Michael Letko, Spyros Lytras, Kei Sato

**Affiliations:** Division of Systems Virology, Department of Microbiology and Immunology, The Institute of Medical Science, The University of Tokyo, Tokyo, Japan; Department of Computational Biology and Medical Sciences, Graduate School of Frontier Sciences, The University of Tokyo, Kashiwa, Japan; Paul G. Allen School for Global Health, Washington State University, Pullman, WA, USA; Department of Epidemiology, Gillings School of Global Public Health, University of North Carolina at Chapel Hill, Chapel Hill, NC, USA; MRC-University of Glasgow Centre for Virus Research, Glasgow, UK; International Research Center for Infectious Diseases, The Institute of Medical Science, The University of Tokyo, Tokyo, Japan; Graduate School of Medicine, The University of Tokyo, Tokyo, Japan; International Vaccine Design Center, The Institute of Medical Science, The University of Tokyo, Tokyo, Japan; Collaboration Unit for Infection, Joint Research Center for Human Retrovirus infection, Kumamoto University, Kumamoto, Japan; Faculty of Medicine, Chulalongkorn University, Bangkok, Thailand; Duke-NUS Medical School, Singapore

**Author notes:** Corresponding authors (Michael Letko), (Spyros Lytras), (Kei Sato). These authors contributed equally (Victoria A. Jefferson, Nicholas J. Catanzaro).

## Abstract

The emergence of novel coronaviruses from animal reservoirs continues to pose a significant zoonotic threat. Here, we investigate the evolutionary origins and virological properties of a recently reported mink-derived HKU5-related coronavirus (nvHKU5r-CoV). Phylogenetic and recombination analyses reveal that nvHKU5r-CoV originated from bat HKU5-like viruses circulating in southeastern China. We characterize the spike loop2 region as a critical determinant of ACE2 receptor specificity, directly interacting with the receptor, and show that the bat HKU5r-CoV with the closest loop2 sequence to nvHKU5r-CoV could already utilize mink ACE2. Using AlphaFold3, we predicted spike-ACE2 binding interfaces consistent with our experimental infectivity results. Targeted mutagenesis demonstrates that a single amino acid substitution (R548S) enables robust entry of nvHKU5r-CoV via human ACE2. We further show that this substitution can arise *in vitro* during hACE2-expressing cell infection with a replication-competent VSV system. Molecular dating suggests that nvHKU5r-CoV transmitted from bats to mink within the last decade, consistent with an expansion of mink fur farming in China. Overall, our findings highlight the zoonotic potential of these viruses and the molecular and structural determinants underlying it, while emphasizing fur farming as a major risk factor for how bat HKU5r-CoVs can transmit to farmed animals and ultimately humans.

## Introduction

The *Merbecovirus* subgenus is a diverse group of coronaviruses^1–3^, several of which pose zoonotic threats and possible emergence^4–6^. The prototype strain that has significant morbidity and mortality in humans is the Middle East respiratory syndrome coronavirus (MERS-CoV), which utilizes dipeptidyl peptidase 4 (DPP4)^7,8^ as its entry receptor and has continuously spilled over into humans from dromedary camels since at least the early 1990’s^9,10^. In contrast, the angiotensin-converting enzyme 2 (ACE2) entry receptor has historically been associated with the entry of sarbecoviruses (subgenus *Sarbecovirus*) including SARS-CoV and SARS-CoV-2^11^; and the seasonal human coronavirus NL63^12^ (subgenus *Setracovirus*). However, recent work has revealed that receptor usage in merbecoviruses is more plastic than previously appreciated^13^. Multiple studies have confirmed that members of the MERS-related coronavirus (MERSr-CoV) group, including NeoCoV, PDF-2180, PnNL2018B and MOW15-22, can utilize ACE2 for host cell entry, revealing that ACE2 usage is not confined to sarbecoviruses and setracoviruses, but also functions as a critical entry receptor across merbecoviruses^14,15^. Merbecoviruses are classified into four main groups based on their phylogenetic relatedness: MERSr-CoVs (*Betacoronavirus cameli*), HKU4-related coronaviruses (HKU4r-CoVs, *Betacoronavirus tylonycteridis*), Hedgehog-CoV-1 *(Betacoronavirus erinacei*), and HKU5-related coronaviruses (HKU5r-CoVs, *Betacoronavirus pipistrelli)*^1,3,6^. While MERS-CoV and HKU4r-CoVs have been known to use DPP4, the receptor usage of HKU5r-CoVs had long remained elusive^16^, partly because of their apparent restricted receptor usage range^14^. Recent studies have solved this conundrum, demonstrating that both HKU5-CoV-1 and HKU5-CoV-2 viruses from *Pipistrellus* species can interact with the ACE2 receptor of their bat hosts^2,6,17^, albeit through different binding conformations than other ACE2-dependent viruses^17^. These findings shed new light on the evolutionary potential of other merbecoviruses like HKU5r-CoVs and their capacity to use different receptor orthologs to mediate cross-species transmission events and zoonoses.

In late 2024, Zhao and colleagues reported the first detection of HKU5r-CoV in farmed mink (*Neogale vison*), an unexpected host, as these viruses have only previously been reported in *Pipistrellus* bats^18^. The farming environment, characterized by dense housing and multispecies contact, creates ideal conditions for viral amplification and interspecies transmission, while fur farming continues to be a major industry in China, involving tens of millions of minks per year^19^. Notably, SARS-CoV-2, another ACE2-dependent coronavirus, has been reported in farmed fur animals and its transmission back into humans has also been observed^20,21^. The discovery of HKU5 viruses in mink positions these animals as a potential intermediate host for HKU5r-CoVs, adding to the growing concerns about fur farms as ecological niches where viral evolution and zoonotic spillover events may be accelerated^22^.

The detection of this HKU5r-CoV in mink prompted investigation into its origin, evolutionary trajectory, and potential for cross-species transmission. To elucidate the emergence of this virus in mink, we performed a comprehensive analysis that integrates phylogenetics, recombination, molecular dating, structural prediction modeling, pseudovirus-based entry assays and full-length recombinant virus replication experiments. Through this integrative approach, we provide molecular and structural insights into how HKU5r-CoVs utilize a merbecovirus-specific loop in their spike receptor binding domain (RBD) to modulate receptor preference and assess the viruses’ spillover potential in light of these findings.

## Results

### Phylogenetic placement of mink HKU5r-CoV

To establish the evolutionary context of the mink-derived HKU5r-CoV (nvHKU5r-CoV), we first conducted BLASTn^23^ searches using the full-length genome and spike gene of nvHKU5r-CoV as queries against the NCBI Genbank database. Through this approach, we retrieved a diverse set of related merbecoviruses, including representatives from MERSr-CoVs, HKU4r-CoVs and HKU5-CoV-1 (44 sequences for the full-length genome, and 61 sequences for the Spike). HKU5-CoV-2 sequences were manually added to the dataset based on recent taxonomic updates^6^.

In the whole genome phylogeny (Extended Data Fig.1a), nvHKU5r-CoV clustered within the HKU5-CoV-1 lineage, without a single closest relative, forming part of a clade that includes several bat-derived viruses from southeastern China. HKU5-CoV-2^6^, described in recent studies as being genetically distinct, is substantially distant to nvHKU5r-CoV and is placed as the sister lineage of the HKU5-CoV-1 clade within the merbecovirus tree. This phylogeny also confirms the broader separation of HKU5r-CoVs from MERSr-CoVs and HKU4r-CoVs. The spike-based phylogeny (Extended Data Fig. 1b) preserved this lineage structure, with HKU5-CoV-1 and HKU5-CoV-2 exhibiting divergent spike gene architectures, further supporting their evolutionary and functional divergence.

To resolve the placement of nvHKU5r-CoV among its closest relatives, we constructed a whole genome phylogeny (Fig. 1a) focusing specifically on HKU5r-CoVs (Table S1), in contrast to the broad merbecovirus tree shown in Extended Data Fig 1a. This tree revealed a clear lineage structure, where three viruses sampled in Zhejiang province (WZ_Pi.abramus_betacorona_2, referred to as WZ2, YJB_Pabr, and BtPa-BetaCoV/ZJ2010-Q280, referred to as Q280) formed a clade basal to nvHKU5r-CoV and its closest relatives. The clade most closely related to nvHKU5r-CoV (which was sampled in the northern province of Shandong) included a number of bat HKU5r-CoV-1 genomes sampled in the southern provinces of Jiangxi, Hainan, and Yunnan. To account for potential confounding effects of recombination within these genomes, we also reconstructed a recombination-free phylogeny by removing recombinant fragments from the whole-genome alignment using RDP5^24^. The resulting tree (Extended Data Fig. 1c) retained the placement of nvHKU5r-CoV close to the Zhejiang bat virus clade, although with reduced bootstrap support, while the number of genomes in the clade closest to nvHKU5r-CoV was largely reduced, indicative of extensive recombination.

**Figure 1.**
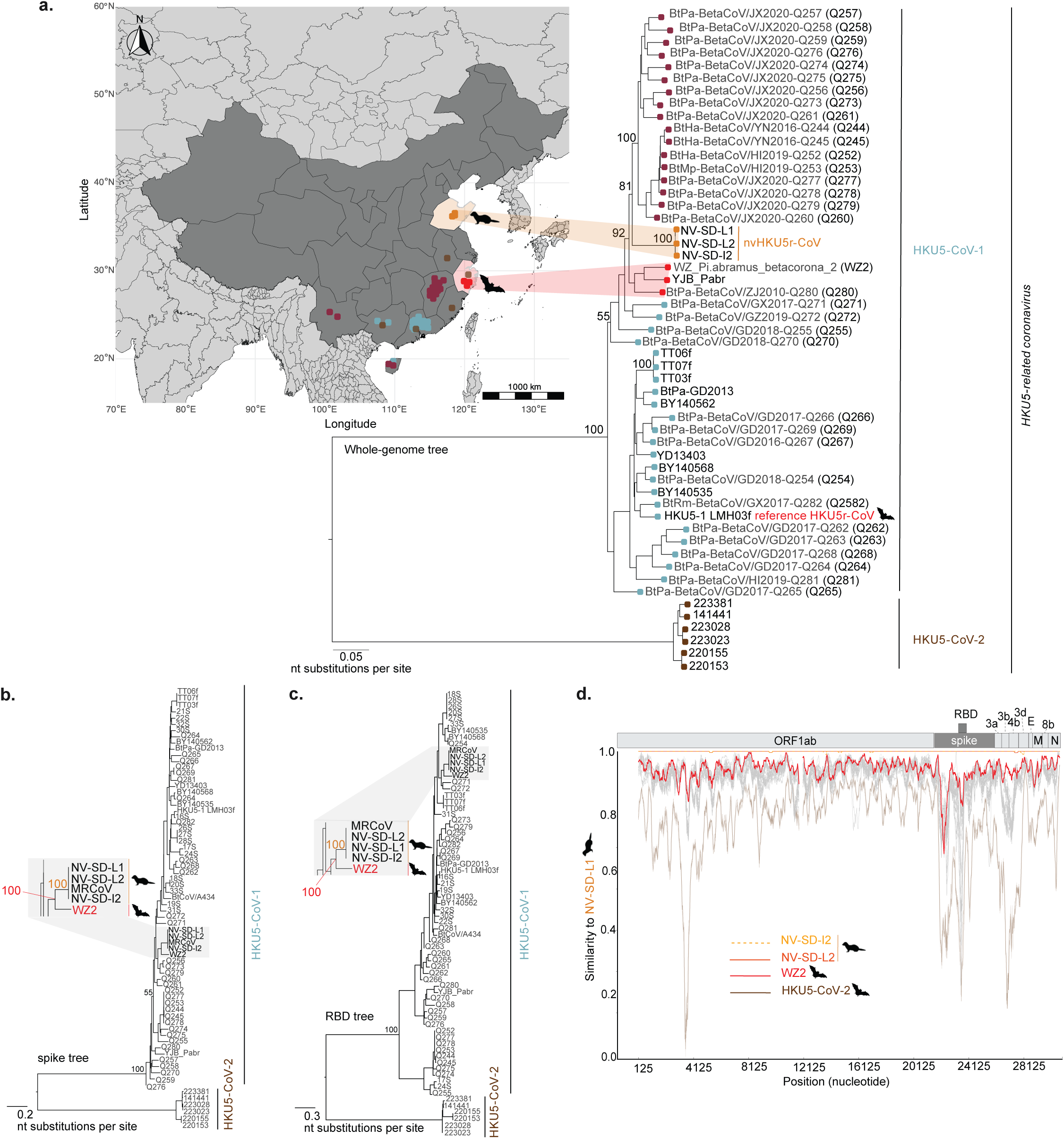
Phylogenetic and genomic characterization of mink-derived HKU5r-CoV. **a.** ML phylogeny based on whole genome sequences of HKU5r-CoVs, showing nvHKU5r-CoV clustering with a lineage of bat-derived viruses from Zhejiang province, including WZ2, YJB_Pabr, and Q280. Downstream lineages include nvHKU5r-CoV from Shandong and other HKU5-CoV-1 viruses from Jiangxi, Hainan, and Yunnan provinces. HKU5-1 LMH03f is highlighted as the reference genome of HKU5r-CoV. **b.** Phylogeny based on full-length spike gene sequences. Zoomed in inset showing nvHKU5r-CoV clustering with WZ2. **c.** Phylogeny based on RBD sequences. Zoomed in inset showing nvHKU5r-CoV clustering with WZ2. Node support for key splits is annotated on each phylogeny. Scale bars represent nucleotide substitutions per site. **d.** Similarity plot analysis using NV-SD-L1 as the reference genome, showing high sequence identity with WZ2 across multiple genomic regions. NV-SD-L1 was selected as the representative nvHKU5r-CoV genome based on >99% identity with NV-SD-I2 and NV-SD-L2.

When looking at phylogenies based on the spike (Fig. 1b) and its RBD (Fig. 1c) we observe that nvHKU5r-CoV confidently clusters most closely with bat virus WZ2, sampled from Zhejiang. This confident topology identifies WZ2 as the single closest known spike sequence to nvHKU5r-CoV’s spike, in contrast to the whole genome phylogeny, which places nvHKU5r-CoV sister to a larger clade of bat viruses.

Similarity plot analysis across the whole genome (Fig. 1d, Extended Data Fig. 1d) further supports this phylogenetic relationship. For comparative analyses, we selected NV-SD-L1 as the representative nvHKU5r-CoV genome due to its good assembly quality and near-complete identity to the other nvHKU5r-CoVs (Extended Data Fig. 1d, 1e). Specifically, NV-SD-L1 and NV-SD-L2 share 99.70% nucleotide identity across the full genome and exhibit 100% identity at both the spike and RBD regions (Extended Data Fig. 1e). For MRCoV, a newly isolated nvHKU5r-CoV from the same mink samples that the NV-SD genomes were sequenced from^25^, only the spike sequence is available, which is identical to that of the other two genomes. The last available genome sequence, NV-SD-I2, is similarly closely related (99.97%-99.98% in the whole genome and identical in the Spike), but has multiple ambiguous bases (‘N’) present in its sequence, particularly within the spike region (Extended Data Fig. 1e). NV-SD-L1 was therefore used as the primary reference for subsequent computational and experimental investigations in this study.

### Evolution of the RBD loop2 region in nvHKU5r-CoV

Given the essential role of the spike protein in mediating host receptor engagement, we compared the RBD sequence of NV-SD-L1 against WZ2 and the HKU5r-CoV reference genome HKU5-1 LMH03f (Fig. 2a). Merbecovirus spike proteins contain two adjacent loop structures in their RBD which directly interact with the entry receptor. The second loop structure (loop2; region 23230-23287 in the NV-SD-L1 genome; 1606-1662 in its spike protein) has been implicated as an essential structural element regulating ACE2 receptor engagement for HKU5r-CoVs^2^. This loop has previously been categorized into short and long structural configurations within HKU5r-CoVs based on indel variation and conformational differences. This suggests that loop2 structural differences may shape host range and cross-species transmission potential by determining the spike – ACE2 orthologs engagement patterns. Therefore, we investigated the evolutionary and structural features of loop2 in nvHKU5r-CoV to understand its role in mink adaptation and zoonotic emergence potential. Our analysis showed that NV-SD-L1 possesses a long loop2 genotype, similar to WZ2, and distinct from the short loop sequence observed in HKU5-1 LMH03f (Fig. 2a). To trace the evolutionary origin of the loop2 region, we first performed a similarity plot analysis across the spike gene (Fig. 2b), which confirmed that most regions of the NV-SD-L1 spike are most similar to WZ2. However, a distinct peak of similarity was observed between NV-SD-L1 and BtPa-BetaCoV/GD2017-Q265 (Q265) specifically within the loop2-encoding region. This coincides with a drop in similarity to WZ2, implying that WZ2 may have recombined away from the nvHKU5r-CoV lineage in this region (Fig. 2a,b). To further investigate this localized similarity, we compared phylogenetic trees constructed from the full RBD and the isolated loop2 region, both rooted using the HKU5-CoV-2 clade as the outgroup (Fig. 2c).

**Figure 2.**
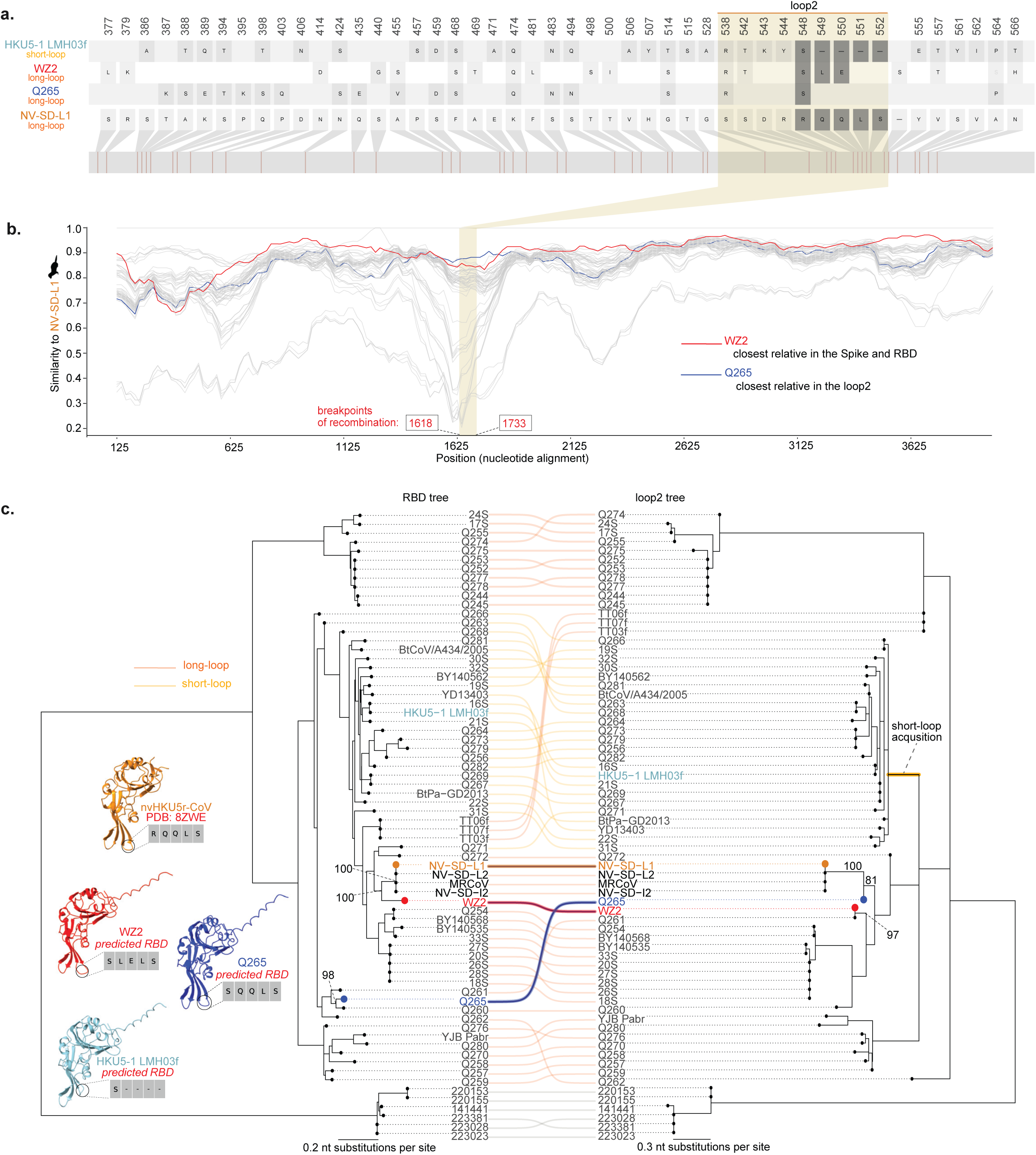
Evolutionary and structural analysis of the spike loop2 region. **a.** RBD amino acid alignment of unique NV-SD-L1 substitutions, highlighting the loop2 region. NV-SD-L1 exhibits a long loop2 configuration (RQQLS), similar to WZ2 (SLELS), and distinct from the short loop structure in HKU5-1 LMH03f (S-----). **b.** Percent identity across the spike gene between NV-SD-L1 and representative HKU5r-CoVs. The GARD-inferred breakpoints around the loop2 region are annotated on the plot. **c.** Tanglegram between phylogenetic trees based on the nucleotide alignment of the RBD-encoding (left) and the loop2-encoding (right) regions. Both trees are rooted by the HKU5-CoV-2 clade. The lines connecting the two trees are coloured by the viruses’ loop2 length (short or long). On the bottom left the following RBD structures are displayed along with their loop2 residues: nvHKU5r-CoV (PDB: 8ZWE), WZ2, Q265, and HKU5-1 LMH03f. Node support for key splits is annotated on each phylogeny. Scale bars represent nucleotide substitutions per site.

The tanglegram comparison revealed a clear topological incongruence: while the RBD tree placed NV-SD-L1 closest to WZ2, the loop2 tree clustered NV-SD-L1 with Q265 (node support = 81), indicative of recombination of this genomic region. Importantly, based on the loop2 tree we can identify a single branch where the deletion that led to the short loop2 genotype took place (Fig. 2c). In contrast, the short loop2 genotype viruses are not monophyletic in the RBD tree, with multiple incongruencies being observed between the two trees. This pattern suggests that multiple recombination events of the loop2 region have likely happened in the evolution of the HKU5r-CoVs.

To investigate this further, we used the genetic algorithm for recombination detection (GARD)^26^ to identify recombination breakpoints within the spike gene. Thirteen breakpoints were detected (Extended Data Fig. 2a), and phylogenies were reconstructed for each resulting non-recombinant region (NRR). In 9 out of the 14 NRRs, WZ2 is the closest or within the clade of closest relatives to the nvHKU5r-CoV (Extended Data Fig. 2b), confirming our results based on the full spike and RBD trees (Fig. 1b,c). However, NRR7 (NV-SD-L1 spike positions 1618–1733) which has breakpoint positions tightly encompassing the loop2 region (Fig. 2b) showed confident phylogenetic clustering between NV-SD-L1 and Q265 (node support = 83), reinforcing the loop2 recombination hypothesis (Extended Data Fig. 2b). This pattern was further supported by examining the genetic distance between NV-SD-L1 and its closest bat virus relative across all NRRs (Extended Data Fig. 2c), where NRR5 and NRR13 exhibited the lowest distances between nvHKU5r-CoV and WZ2 (0.0151 and 0.0215 substitutions per site, respectively). Together, these results confirm that WZ2 has the overall closest spike sequence to nvHKU5r-CoV, with the exception of the loop2 region, where Q265 is the closest known relative. The phylogenetic analyses suggest that this is likely a result of both Q265 acquiring this closest loop2 through recombination, as well as WZ2 acquiring a slightly more distant loop2 segment.

To further characterize the structural implications of loop2 variation, we examined the RBD structure of nvHKU5r-CoV using the MRCoV isolate (PDB: 8ZWE)^25^ as the reference. MRCoV shares 99.98% identity at the spike gene and 100% identity in the RBD sequence with NV-SD-L1 (Fig. 1b-c, Extended Data Fig. 1e), and its structure has already been determined using cryo-EM^25^, making it a reliable model for nvHKU5r-CoV. In this structure, the loop2 sequence of nvHKU5r-CoV, RQQLS, was identified. We then predicted the RBD structures of WZ2 (SLELS), Q265 (SQQLS), and HKU5-1 LMH03f (S----, representing a short loop) using AlphaFold3 (AF3)^27^. These comparisons revealed notable structural diversity in the loop2 region across HKU5r-CoVs. In the cryo-EM structure of MRCoV (representing NV-SD-L1), the loop2 adopts an extended conformation with the RQQLS motif. The predicted structure of Q265, which shares the most similar loop2 sequence (SQQLS) to NV-SD-L1, also displays an extended loop, closely resembling that of nvHKU5r-CoV. WZ2 encodes a slightly different loop2 motif (SLELS) but still has a long loop genotype. In contrast, HKU5-1 LMH03f, representing the short loop variant, lacks the extended insertion entirely, resulting in a more compact loop2 structure.

These structural differences suggest that loop2 extension may influence the accessibility and orientation of key residues involved in ACE2 binding. Long loop configurations, such as those in NV-SD-L1, Q265, and WZ2, may enhance receptor engagement by projecting interaction residues outward, while short loop variants may restrict binding due to steric limitations. This interpretation is supported by our recent work^2^ demonstrating that short loop architectures can constrain RBD orientation and limit ACE2 compatibility in certain HKU5r-CoVs.

### Functional and structural determinants of ACE2 usage in HKU5r-CoVs

To assess the receptor usage of HKU5r-CoVs including nvHKU5r-CoV, we performed pseudovirus entry assays using cells expressing ACE2 orthologs from *Neogale vison* (nvACE2) and humans (hACE2) (Fig. 3a). The long loop2 NV-SD-L1 RBD exhibited robust entry via nvACE2, consistent with it being sampled in mink^18,25^. Notably, several short loop HKU5r-CoVs also mediated entry via nvACE2. In contrast, the majority of long loop2 HKU5r-CoVs failed to enter nvACE2-expressing cells, with the surprising exception of the Q265 RBD. As discussed above, Q265 has the closest loop2 region to the nvHKU5r-CoVs (Fig. 2), suggesting that i) loop2 architecture plays a key role in determining receptor compatibility, and ii) long loop2 HKU5r-CoVs with similar loop2 sequences circulating in bats are capable of using mink ACE2 for cell entry. Consistent with previous research, only the HKU5-CoV-2-441 RBD was able to mediate weak entry via hACE2^6^ indicating a general lack of compatibility of all other known HKU5r-CoVs with the human receptor.

**Figure 3.**
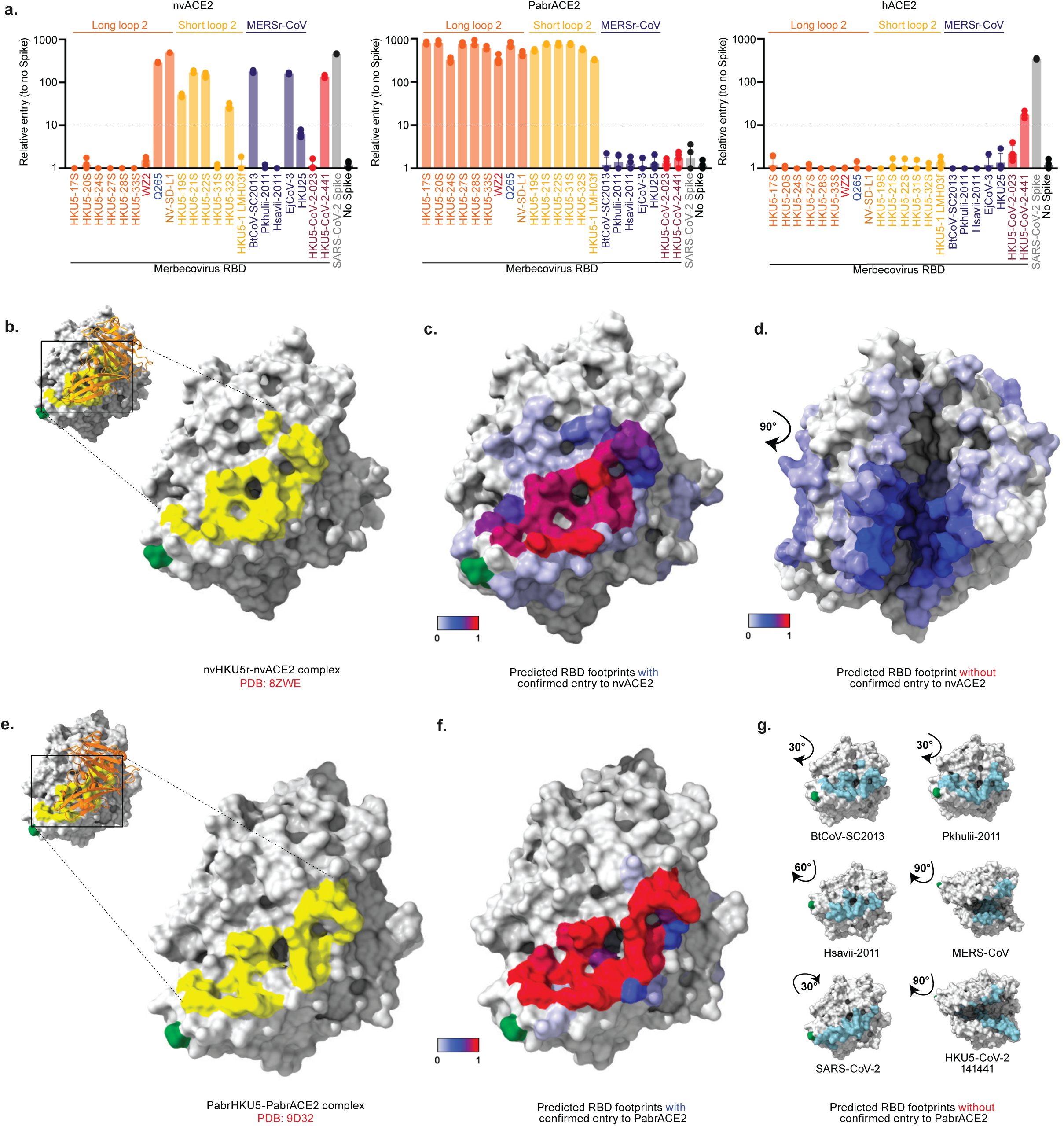
Functional and structural determinants of ACE2 usage in HKU5r-CoVs. **a.** Pseudovirus entry assays using Vero cells transduced to stably express ACE2 orthologs from *Neogale vison* (nvACE2), *Pipistrellus abramus* (PabrACE2) and human (hACE2). nvHKU5r-CoV, Q265, and short loop2 HKU5r-CoVs efficiently entered nvACE2-expressing cells, while other long loop2 viruses failed to mediate entry. All HKU5r-CoV entered via PabrACE2. SARS-CoV-2 served as a positive control. The dotted line indicates the assay’s detection limit at 10 relative entry units. Four technical replicates are displayed on each bar. Binding footprint or proportion of footprint interactions of spike for: **b.** Native nvHKU5r-CoV RBD bound to nvACE2 (PDB: 8ZWE), **c.** AF3-predicted models of entry-positive HKU5r-CoVs on nvACE2, **d.** AF3-predicted models of entry-negative HKU5r-CoVs on nvACE2, **e.** Native PabrHKU5r-CoV–PabrACE2 complex (PDB: 9D32), **f.** AF3-predicted models of entry-positive HKU5r-CoVs on PabrACE2, **g.** AF3-predicted of entry-negative merbecoviruses and SARS-CoV-2 on PabrACE2.

To further explore receptor compatibility across species, we integrated structural modeling of spike–ACE2 interactions using AF3^27^, that includes prior experimental data from *Pipistrellus abramus* ACE2 (PabrACE2), which was tested in our earlier work^2^ (Table S2). Cryo-EM co-structures of nvHKU5r-CoV and nvACE2 (PDB: 8ZWE)^25^, and PabrHKU5r-CoV and PabrACE2 (PDB: 9D32)^17^ served as references for evaluating predicted co-structures. To understand how different HKU5r-CoVs interact with ACE2 receptors, we examined the structural footprints formed at the interface between RBD and ACE2. These footprints represent specific contact points where the viral RBD physically interacts with ACE2.

To visualize how different HKU5r-CoVs and other tested viruses interact with ACE2 receptors, we compared reference cryo-EM structures and AF3-predicted models of RBD-ACE2 complexes across nvACE2 and PabrACE2 (Fig. 3b-g). The cryo-EM structure of the nvHKU5r-CoV RBD bound to nvACE2 (Fig. 3b) reveals a compact and surface-accessible RBD footprint, which closely resembled predicted structures of pseudovirus entry-positive RBDs, including HKU5r-CoVs and other merbecoviruses (Fig. 3c). Consistent with our experimental results, the four short loop2 RBDs that infected nvACE2 (19S, 21S, 22S, and 32S), Q265 and NV-SD-L1 had confident predictions (ipTM and pTM equal or above 0.80) interacting with the same part of nvACE2. In contrast, pseudovirus entry-negative RBDs exhibit dispersed and misaligned contacts on nvACE2 (Fig. 3d) with low prediction confidence, the only exception being HKU5-33S which had a confident prediction consistent with entry-positive RBDs but showed no entry on the pseudovirus assay (Fig. 3a). For the bat ACE2, the native PabrHKU5r-PabrACE2 structure (Fig. 3e) and predicted models of entry-positive RBDs (Fig. 3f) again show well-aligned RBD footprints. These structural comparisons highlight that AF3-based HKU5r-CoV RBD binding predictions to different ACE2 orthologs are generally consistent with experimental pseudovirus results and predictions between non-interacting molecules are likely inaccurate.

Using these predicted co-structures, we can identify the specific residues utilized by the HKU5r-CoVs for binding each of the three tested ACE2 residues. In nvACE2, the key contact residues were Glu30, Tyr34, Glu37, Tyr41, Lys353, His354, Asp355, and Arg393. PabrACE2 had a wider consistent footprint composed of Arg26, Val30, Asn33, His34, Asp90, Ile93, Gln96, Met322, Thr323, Pro324, Trp327, Arg328, Lys352, Asn353, Ala385, Asn386, and Ser388, although it should be noted that the larger number of contact sites may be a result of more RBDs successfully binding PabrACE2. In contrast, entry-negative RBDs exhibited dispersed and inconsistent footprints, often targeting internal or sterically hindered regions of ACE2, suggesting structurally incompatible receptor engagement (Extended Data Figs. 3–5).

Still, some predicted co-structures deviated from the expected patterns. For instance, Pkhulii-2011^28^ detected in *Pipistrellus kuhlii* and Hsavi-2011^28^ identified from *Hypsugo savii* were predicted by AF3 to engage nvACE2 and PabrACE2 (Extended Data Figs. 3, 4), despite being entry-negative in pseudovirus assays. Their predicted RBD-ACE2 interfaces showed variable confidence (Table S2), for nvACE2, Pkhulii-2011 had a pTM/ipTM values of 0.66/0.15, while Hsavi-2011 scored 0.74/0.64; for PabrACE2, Pkhulii-2011 reached 0.74/0.60, and Hsavi-2011 again showed low interaction site confidence at 0.65/0.15. These values suggest that while the RBD and ACE2 monomer structures were reasonably well predicted, the binding interaction between the two proteins had poor confidence prediction. This discrepancy may reflect lineage-specific incompatibilities, as both viruses belong to the MERSr-CoV group rather than the HKU5r-CoV lineage identified in *P. abramus*. This was also the case with HKU5-CoV-2-441, which was predicted to have no significant binding to any tested ACE2 (ipTM below 0.30 for nvACE2, PabrACE2, and hACE2), even though it can use both mink and, weakly, human ACE2 based on our infectivity assays (Fig. 3a)^6^.

To benchmark the accuracy of our structural predictions, we compared AF3 predictions with those from Boltz-1^29^ and chai-1^30^. AF3 consistently produced high-confidence models with pTM and ipTM values supporting the reference cryo-EM conformations (Extended Data Fig. 6a). In contrast, the two alternative methods yielded lower confidence scores and failed to resolve key interface features, particularly in the loop2 regions (Extended Data Fig. 6b-c).

### Replication of nvHKU5r-CoV in cells expressing mink ACE2

To validate our pseudovirus infectivity results, we further assessed receptor usage and viral replication of full genome HKU5r-CoVs: NV-SD-L1 and HKU5-5 (both long loop2). Recombinantly derived reporter viruses expressing nano-luciferase (nLuc) (Fig. 4a) were evaluated in Vero81 cells expressing human, mink, *P. abramus*, or mouse ACE2 (mACE2), with empty vector cells as a negative control. HKU5-5 mediated strong entry through PabrACE2, with only weak entry in nvACE2, and no detectable entry in mACE2 or hACE2 (Fig. 4b). In contrast, NV-SD-L1 entered efficiently using both PabrACE2 and nvACE2 at near equal levels but again showed no entry with either mACE2 or hACE2.

**Figure 4.**
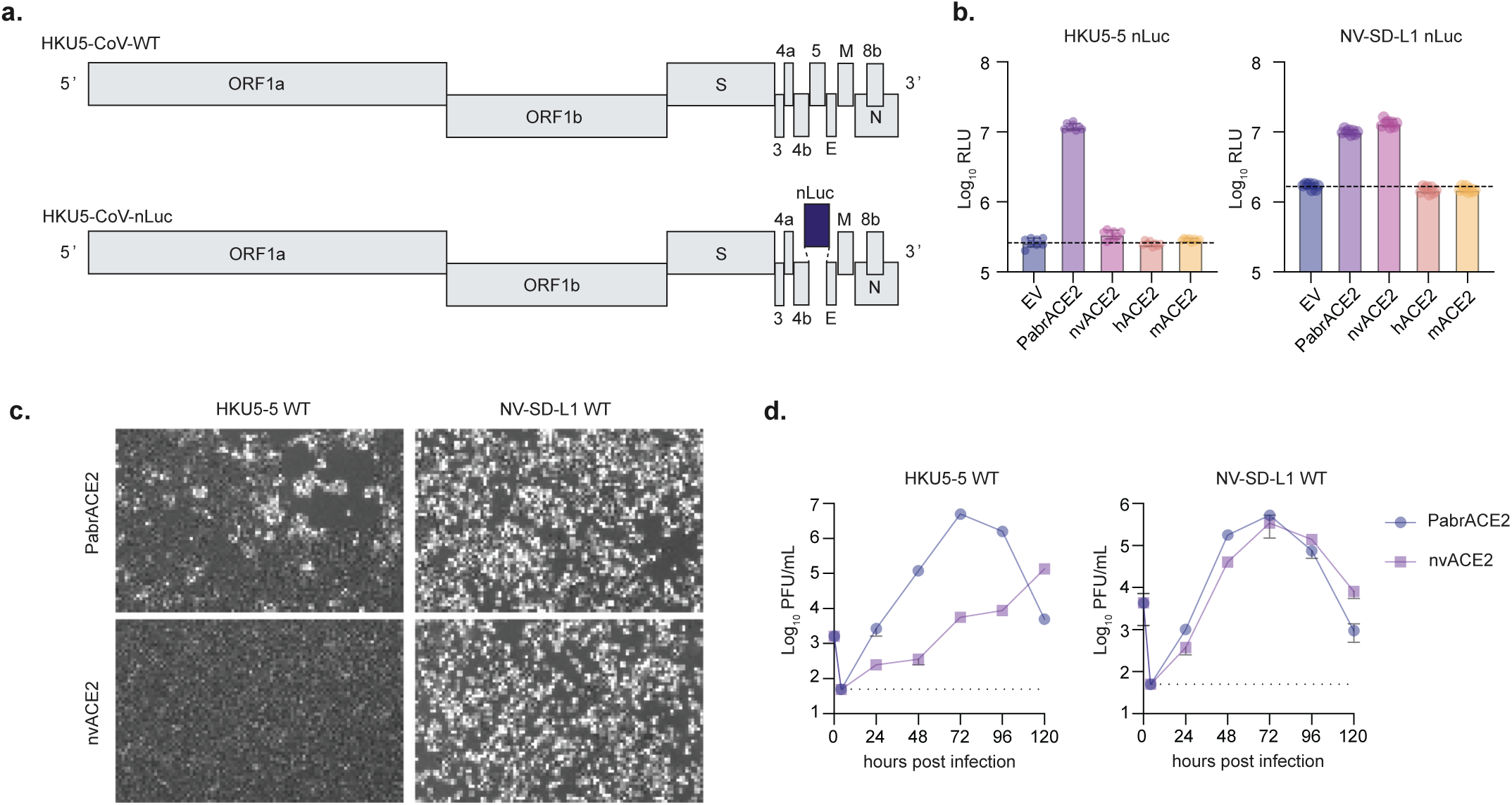
*In vitro* replication of HKU5r-CoVs in cells expressing different ACE2 orthologs. **a.** Schematic of full-length recombinant HKU5r-CoV constructs encoding wild-type or nano-luciferase (nLuc) reporters for HKU5-5 (long loop2) and NV-SD-L1 (mink-derived, long loop2). **b.** Entry efficiency of HKU5-5 and NV-SD-L1 nLuc viruses in Vero81 cells expressing ACE2 orthologs from mink (nvACE2), *Pipistrellus abramus* (PabrACE2), human (hACE2), or mouse (mACE2). Data are normalized to empty vector (EV) controls. The dotted line indicates the mean EV control RLU value. Eight technical replicates are displayed on each bar. **c.** Representative cytopathic effect (CPE) in nvACE2- and PabrACE2-expressing cells infected with HKU5-5 or NV-SD-L1. **d.** Multi-step growth curves of HKU5-5 and NV-SD-L1 in nvACE2 or PabrACE2 expressing Vero81 cells following infection at an MOI of 0.01. Viral titers were quantified at the indicated time points by plaque assay. The dotted lines indicate the assay’s detection limit.

To determine whether this entry translated into productive infection, we performed multi-step growth curves in susceptible cells. Consistent with our entry assays, NV-SD-L1 exhibited robust replication in cells expressing PabrACE2 or nvACE2, confirming its adaptation to this host receptor (Fig. 4c,d). HKU5-5 showed robust replication in PabrACE2 cells, and detectable but delayed and low-level replication in nvACE2 cells (Fig. 4c,d). This is supported by AF3 structural predictions, which indicated a compatible RBD footprint on nvACE2 (Extended Data Fig. 8). These results demonstrate the plasticity that HKU5r-CoV RBDs have in receptor specificity. Consistent with NV-SD-L1, it seems that some long loop2 RBDs have the potential to enable entry via nvACE2 and allow viral replication.

### Loop2-targeted mutagenesis reveals key determinants of mink ACE2 compatibility for HKU5r-CoVs

To further investigate the molecular basis of ACE2 receptor specificity among HKU5r-CoVs, we focused on the loop2 region. Based on our results, the NV-SD-L1, Q265 and short loop2 HKU5r-CoV RBDs mediate entry via nvACE2 and PabrACE2, whereas all other tested long loop2 viruses fail to enter cells via nvACE2. This phenotypic discrepancy suggests that loop2 length alone does not determine receptor usage, and that specific residues within the loop may modulate receptor compatibility, as also shown by our virus replication results (Fig. 4). To explore which specific sites in the RBD may control this phenotype, we performed targeted mutagenesis guided by two approaches: i) a comparative sequence analysis in which we identified consensus residues separately for short and long loop2 viruses and then contrasted these with the sequence of NV-SD-L1 to pinpoint correlated sites (Extended Data Fig. 7a); and ii) structural modeling of loop2 electrostatic surface potential, which allowed us to evaluate how changes in charge distribution across the loop2 region might influence receptor compatibility (Extended Data Fig. 7b).

A deletion after NV-SD-L1 site 554 and a glycine instead of a serine at NV-SD-L1 site 567, were identified to be shared between short loop2 viruses and NV-SD-L1 (Fig. 5a). These changes were introduced into HKU5-20S to mimic the loop2 genotype of the short loop and NV-SD-L1 RBDs. In nvACE2-expressing cells, neither mutation restored entry, indicating that these changes alone are insufficient to mimic the receptor engagement profile of HKU5-21S or NV-SD-L1. The site 554 deletion reduced entry efficiency to PabrACE2, while the S567G substitution had no effect compared to wild-type HKU5-20S.

**Figure 5.**
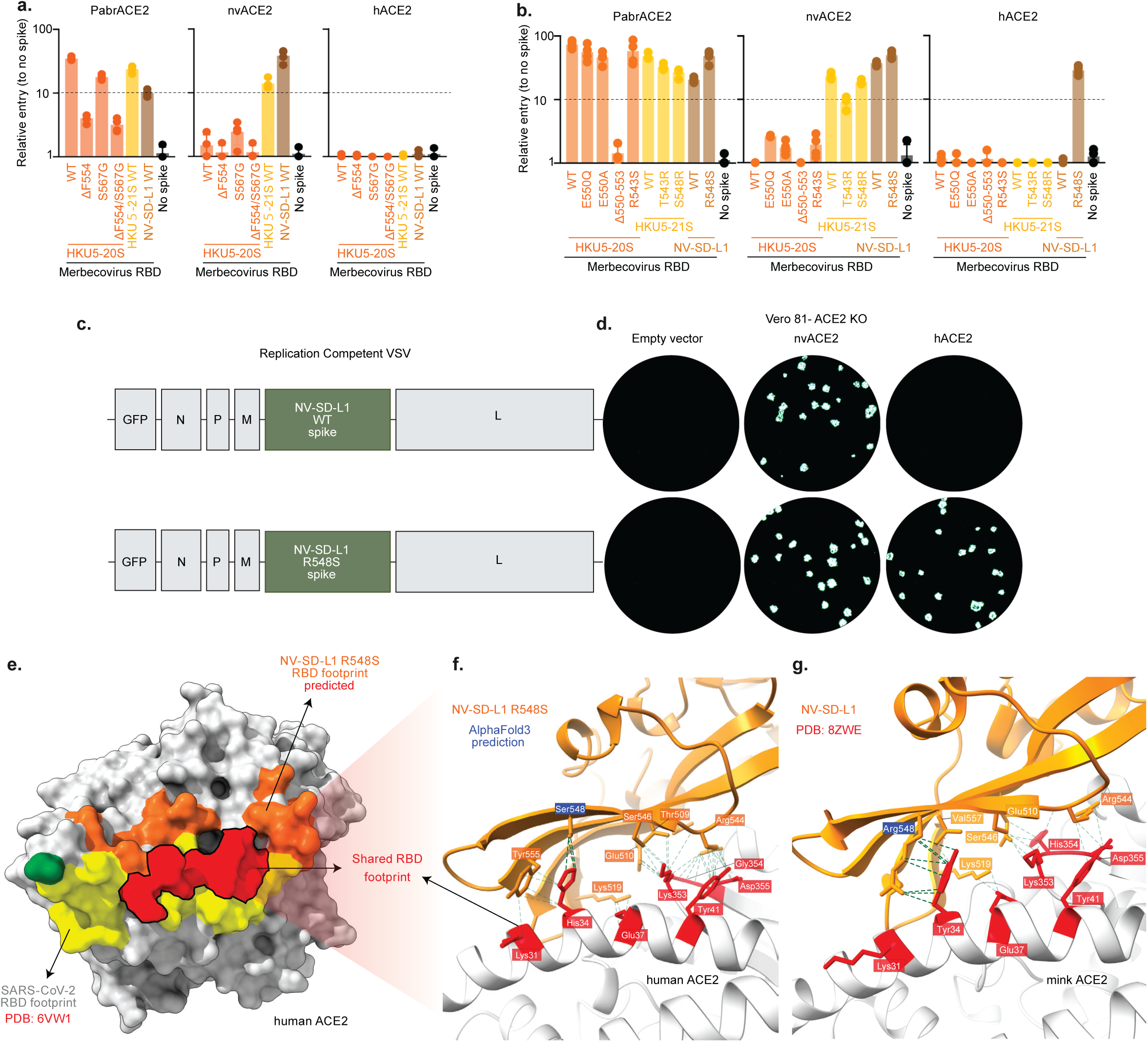
Loop2-targeted mutagenesis reveals determinants of ACE2 compatibility in HKU5r-CoVs. **a.** Pseudovirus entry assays testing loop2 mutations based on correlated sequence sites. **b.** Pseudovirus entry assays testing loop2 mutations identified to contribute to electrostatic interactions between spike and ACE2. The dotted line indicates the assay’s detection limit at 10 relative entry units. Four technical replicates are displayed on each bar. **c.** Schematic of rcVSV constructs encoding the NV-SD-L1 spike of either WT or R548S mutant. **d.** Fluorescent focus-forming assay showing entry of rcVSV (WT or R548S) into Vero81 cells stably expressing nvACE2 or hACE2, or empty vector control. Representative GFP images at 24 hpi. **e.** Structural modeling of the NV-SD-L1 R548S mutant RBD footprint on hACE2, compared to the SARS-CoV-2 RBD footprint (PDB: 6VW1). Conserved hACE2 residues engaged by both viruses are highlighted in red. **f.** Close-up view of the NV-SD-L1 R548S – hACE2 interface showing that Ser548 forms a contact with His34 of hACE2. **g.** Comparison with the native nvHKU5r-CoV–nvACE2 complex (PDB: 8ZWE), where Arg548 in the mink-adapted virus engages Tyr34 of nvACE2.

Electrostatic potential of the loop2 interface exhibited notable variation within long loop2 RBDs. For example, HKU5-20S possessed a bulkier, more negatively charged loop2 compared to NV-SD-L1, potentially impeding nvACE2 binding. Substitution of glutamate at position 550 with glutamine (E550Q, present in NV-SD-L1) or alanine (E550A) slightly enhanced nvACE2-mediated entry, supporting the hypothesis that charge reduction or mimicry of the NV-SD-L1 loop conformation improves compatibility (Fig. 5b). However, deletion of residues Q550-P553 in HKU5-20S (549–552 in NV-SD-L1) to mimic the short loop2 genotype of HKU5-21S abolished entry via PabrACE2 and failed to restore nvACE2 entry (Fig. 5b), suggesting that loop truncation alone is insufficient and may disrupt broader structural integrity.

At position 543 (542 in NV-SD-L1), HKU5-20S encodes an arginine that interacts favorably with a negatively charged pocket in PabrACE2 (D49), but clashes with the positively charged R49 in nvACE2. Substitution to serine (R543S), as found in NV-SD-L1, increased nvACE2 entry. Conversely, introducing an arginine at this position in HKU5-21S (T543R) reduced nvACE2 entry and may have modestly impaired PabrACE2 entry, indicating a context-dependent effect of charge and side-chain volume at this site. It should be noted that Q265 also has a serine at this site (Fig. 2a), partly explaining its ability to use nvACE2, similar to NV-SD-L1.

### A single mutation enables nvHKU5r-CoV to use human ACE2

The last site we identified using our mutagenesis approach is NV-SD-L1 position 548, where NV-SD-L1 encodes an arginine that is predicted to engage Y34 in nvACE2. This residue is directly upstream the loop2 deletion leading to the short loop genotype and arginine is unique to the nvHKU5r-CoV RBD (Fig. 2c). Replacing this residue with serine (R548S), as found in HKU5-20S, HKU5-21S, and Q265 (among other HKU5r-CoVs) increased PabrACE2 entry, while the reciprocal mutation in HKU5-21S (S548R) decreased PabrACE2 entry (Fig. 5b). Perhaps more surprisingly, the R548S substitution on NV-SD-L1 resulted in robust entry in human ACE2 expressing cells (∼90% relative to NV-SD-L1 wild-type; Fig. 5b). In contrast to the inability of all wild-type HKU5r-CoV RBDs to use hACE2 (with the exception of HKU5-CoV-2-441) (Fig. 3a), a single mutation seems to make the nvHKU5r-CoV RBD compatible to hACE2. In addition, nearly all other HKU5r-CoVs have a serine at their corresponding 548 sites, suggesting that this residues effect is further determined by the rest of the NV-SD-L1 RBD.

To explore the potential of this mutation arising on the NV-SD-L1 spike, we employed an *in vitro* evolution approach to identify human receptor tropism determinants using replication-competent VSV (rcVSV) encoding the spike of NV-SD-L1 (Fig. 5c). Mixed cell populations containing 10% PabrACE2 and 90% hACE2 Vero81 cells were infected and serially passaged. By 4 days post-infection, strong GFP signal was observed in all cells in this mixed population. Virus harvested from these co-cultures were plaque purified and sequenced. Strikingly, the R548S was consistently identified in the sequenced virus. When re-introduced into the rcVSV clone, the R548S mutation was both necessary and sufficient to facilitate entry into hACE2-expressing cells as evidenced by spread of GFP positive foci in Vero81 cells expressing nvACE2 and hACE2, but not the empty vector controls (Fig. 5d).

Structural modeling of the NV-SD-L1 R548S mutant RBD binding to nvACE2 (Fig. 5f) revealed partial overlap with the experimentally determined SARS-CoV-2 RBD footprint on hACE2 (PDB: 6VW1; Fig. 5e), while the wild-type NV-SD-L1 RBD exhibited sterically hindered and misaligned interface with no apparent binding to hACE2, consistent with experimental result (Extended Data Fig. 9). The predicted RBD footprint of the NV-SD-L1 R548S mutant and the experimentally determined SARS-CoV-2 RBD footprint both engaged a set of conserved hACE2 residues consisting of Lys31, His34, Glu37, Tyr41, Lys353, Gly354, and Asp355 (Table S3), which are known to stabilize RBD binding and facilitate efficient entry^25,31^. The serine residue on site 548 is predicted to directly interact with the histidine at hACE2 site 34. The 34H hACE2 residue interacts in turn with NV-SD-L1 site 555Y which further stabilizes engagement by contacting hACE2 site 31 (Fig. 5f). Notably, both the PabrACE2 and the hACE2 have a histidine at site 34, while nvACE2 has a tyrosine. In the case of nvACE2 interacting with the wild-type NV-SD-L1 RBD, the arginine at site 548 fully engages the tyrosine at site 34, without the need for spike site 555 and ACE2 site 31 to be involved in the interaction (Fig. 5g). Since both arginine and histidine are positively charged residues R548 is expected to deter the interaction with H34, suggesting that S548R is a mink-specific adaptation acquired to improve binding with the Y34 unique to mink ACE2.

Overall, these findings suggest that while the currently known HKU5r-CoVs are generally incompatible with hACE2, single and naturally occurring mutations like R548S can occur during NV-SD-L1 infection and unlock a critical molecular shift toward human receptor adaptation, enable by direct binding interactions between the receptor and the virus RBD.

### Molecular dating of the emergence of HKU5r-CoV from bats to mink

A remaining question is how much time nvHKU5r-CoV had to acquire its unique features and emerge in mink after it diverged from the closest known bat HKU5r-CoV. To investigate this timescale, we performed molecular dating analyses across the entire genome of the viruses using NRRs inferred by GARD. A total of 46 recombination breakpoints were identified in the complete genome (Fig. 6a), enabling the partitioning of the alignment into 47 NRRs. These regions varied in length from 204 to 2145 nucleotides (nt), with a median of 567 nt, and were analyzed independently using a relaxed molecular clock model (Table S4). Focusing on each NRR, we explored when the closest-inferred viruses ancestral to nvHKU5r-CoV circulated. These are defined as the most recent common ancestor (MRCA) between nvHKU5r-CoV and its closest bat virus relatives.

**Figure 6.**
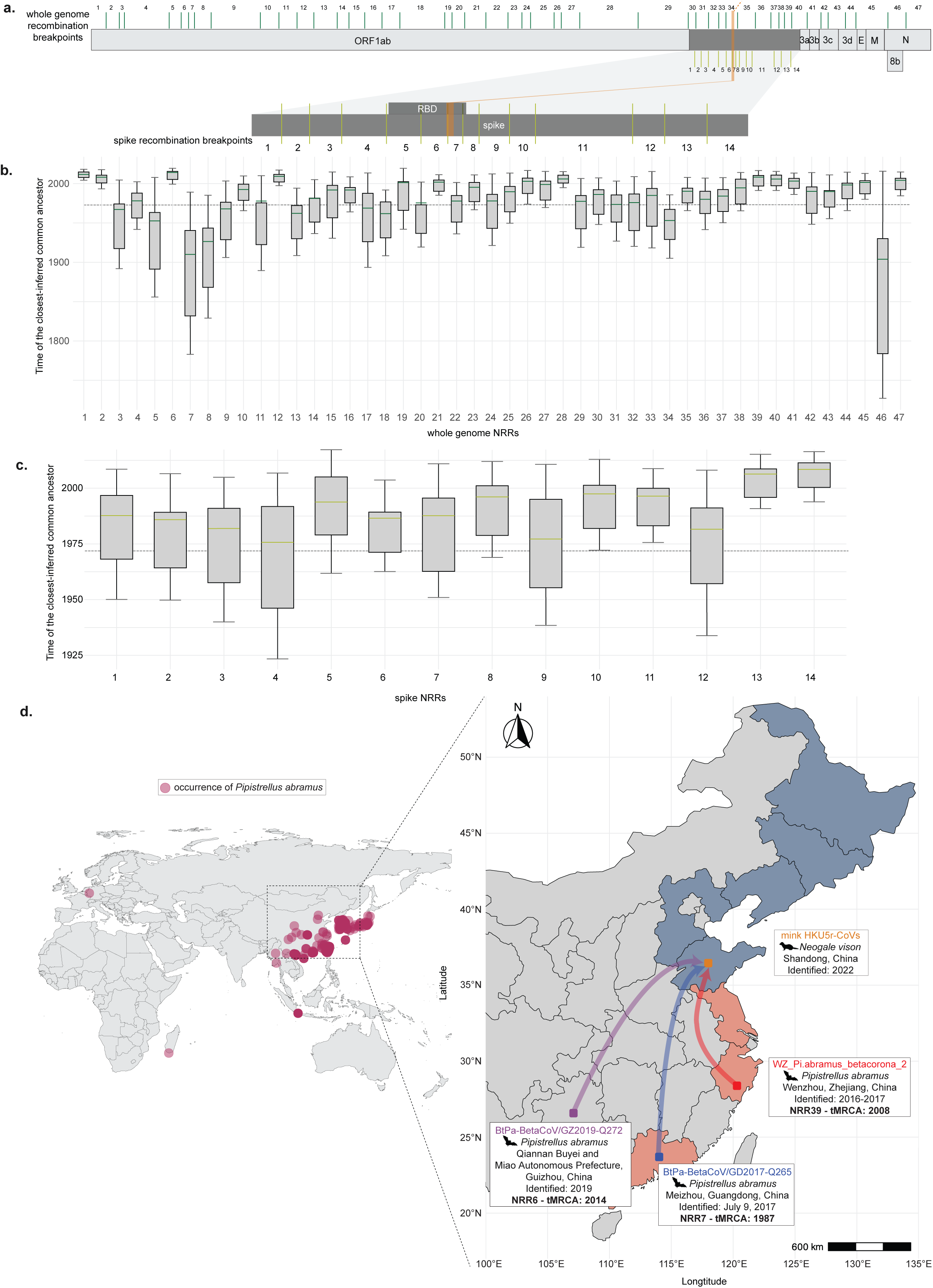
Molecular dating and geographic distribution of nvHKU5r-CoVs closest-inferred ancestors and relatives. **a.** Genome schematic mapping the recombination breakpoint coordinates identified by the GARD whole genome analysis (top). A total of 46 breakpoints were detected genome-wide, allowing for partitioning into 47 NRRs. Spike gene schematic mapping the recombination breakpoint coordinates identified by the GARD spike-specific analysis (bottom). A total of 13 recombination breakpoints, allowing for partitioning into 14 NRRs. The position of the loop2-encoding region is highlighted in both schematics. **b.** Distribution of inferred times to the MRCA between nvHKU5r-CoV and its closest bat virus relatives across all 47 whole-genome NRRs. **c.** Distribution of inferred times to the MRCA between nvHKU5r-CoV and its closest bat virus relatives across all 14 spike NRRs. The dotted lines correspond to 50 years before the emergence of nvHKU5r-CoV in both panels B and C. **d.** Global distribution of *Pipistrellus abramus*, the reservoir host of HKU5r-CoVs, based on the GBIF database (left). Sampling locations of: nvHKU5r-CoV in *Neogale vison* from Shandong (2022), WZ2 from Wenzhou, Zhejiang (2016–2017; tMRCA of 2008), Q265 from Guangdong (2017; tMRCA of 2008), and Q272 from Guizhou (2019; tMRCA of 2014) (right). Chinese provinces shaded in blue indicate major fur farming regions, while Chinese provinces shaded in pink represent primary fur processing and wholesale hubs^19^.

Across all NRRs (Fig. 6b), the inferred times of the most recent common ancestors (tMRCA) ranged between 1903 and 2014, with a median tMRCA of 1989 (Table S4). The two NRRs with the most recent closest-inferred ancestor of nvHKU5r-CoV had an inferred date of 2014 (95% highest posterior density [HPD]: 1999-2019; NRR6 [ORF1ab]; 269nt) and 2011 (95% HPD: 2004-2018; NRR1 [ORF1ab]; 525nt), 8 and 11 years prior to its detection in mink in 2022^18^. The genome most closely related to nvHKU5r-CoV within NRR6 is Q272, sampled in Guizhou, China, whereas the genomes most closely related to nvHKU5r-CoV within NRR1 are Q277, Q278 and Q279, all sampled from Jiangxi, China. Importantly, NRR28, the longest region (2145nt) yielded a tMRCA of 2005 (95% HPD: 1994-2015), 16 years before the detection of HKU5r-CoV in minks. Within NRR28, the single genome most closely related to nvHKU5r-CoV is Q259 sampled in Jiangxi, China. The oldest time of the closest-inferred bat virus ancestors within a single NRR of nvHKU5r-CoV was 1903 (95% HPD: 1994-2015, NRR46 [ORF8b]). Only 10 NRRs (NRRs 3, 5, 7, 8, 9, 13, 17,18, 34, and 46) have a median tMRCA older than 50 years prior to the time the nvHKU5r-CoVs were sampled. Examining the closest-inferred ancestors in all whole genome NRRs, WZ2, sampled from Wenzhou, Zhejiang Province in China, was identified as the closest bat virus relative in 16 out of 47 NRRs (Table S4), making it the closest inferred bat virus ancestor of nvHKU5r-CoV for most of its genome.

To account for finer signals of recombination within the spike protein, we performed molecular dating across the 14 NRRs identified specifically using the spike gene (Fig. 6c, Table S4). These spike NRRs revealed tightly clustered recombination breakpoints (Extended Data Fig. 2), distinct from the broader patterns observed in the whole-genome analysis (Fig. 6a). The inferred median times of the tMRCA for nvHKU5r-CoV across these regions ranged from 1975-2008 with a median of 1987 (Table S4). Among these 14 spike NRRs (Fig. 6a,c), the longest region was spike-NRR11 (807nt), which yielded a median tMRCA of 1996 (95% HPD: 1975-2008), 26 years before the detection of HKU5r-CoV in minks. Within spike-NRR11, the single genome most closely related to nvHKU5r-CoV is WZ2. The spike NRR with the most recent tMRCA to nvHKU5r-CoV were spike-NRR14, with a median tMRCA of 2008 (95% HPD: 1993-2016), 14 years prior to its detection in minks in 2022^18^. The genome most closely related to nvHKU5r-CoV within spike-NRR14 is Q259, sampled in Jiangxi, China. The oldest time of the closest-inferred bat virus ancestors within a single spike-NRR of nvHKU5r-CoV was 1975 (95% HPD: 1923-2006). Interestingly, the median tMRCA of each spike NRR was more recent than 50 years prior to the time the nvHKU5r-CoVs were sampled.

Molecular dating of the spike NRRs where WZ2 is the closest bat relative reveals a consistent evolutionary signal. Among the 14 spike NRRs analyzed, WZ2 was identified as the closest bat ancestor in 9 regions: spike-NRR1, 4, 5, 6, 8, 10, 11, 12, and 13. These regions yielded median tMRCA estimates ranging from 1975 (95% HPD: 1923-2006) to 2006 (95% HPD: 1990-2015). This pattern mirrors the whole-genome analysis (Fig. 6b), where WZ2 was also identified as the closest inferred bat ancestor across most of the genome, further supporting its central position in the evolutionary trajectory of nvHKU5r-CoV. Assessing the date of spike loop2 region, captured by spike-NRR7, reveals a median tMRCA of 1987 (95% HPD: 1950-2010). This region spans 125 nucleotides and was inferred to have Q265 as its closest bat virus relative. Notably, its estimated tMRCA aligns well with the overall median tMRCA of all Spike NRRs.

To contextualize the emergence of nvHKU5r-CoV, we examined the geographic distribution of this virus and its closest bat virus relatives (Fig. 6d). *Pipistrellus abramus*, the natural host of HKU5r-CoVs is primarily found in south and eastern China and Japan^32^.

Consistent with the bats’ distribution, nvHKU5r-CoV, representing the first detection of these viruses in a non-bat host, was identified in a mink farm located in Shandong province^18^. Alongside this virus, Q272, which exhibited the most recent tMRCA to nvHKU5r-CoV (NRR6), was sampled in 2019 from the Qiannan Buyei and Miao Autonomous Prefecture in Guizhou^33^. Q265, the closest relative based on the loop2-encoding region, which can also use nvACE2 (Fig. 3a), was collected in July 2017 from Menzhou in Guangdong^33^. Lastly, WZ2, the virus with the highest overall genomic similarity to nvHKU5r-CoV across the full genome (Fig. 1a), spike gene (Fig. 1b), and the majority of NRRs (Fig. 6a, Extended Data Fig. 2, Table S4) was sampled between 2016 and 2017 in Wenzhou, Zhejiang^34^. This data suggests that nvHKU5r-CoV transmitted to mink within the last decade from a bat HKU5r-CoV lineage likely circulating in the southeastern China, eventually moving to the northeastern Shandong Province where mink fur farming is common^19^.

## Discussion

Mink farming is a global multi-billion-dollar industry which remains largely active in China. Our molecular dating inference of nvHKU5r-CoV’s proximal ancestor having transmitted to mink within a decade prior to the virus’s detection in 2022 is in line with a substantial increase in mink pelt production recorded in China in 2014^19^, increasing the risk of bat-to-mink virus transmission and successful onward transmission within the expanding farmed mink population. In the present study we take a integrative approach, combining phylogenetics, recombination analysis, structural modeling, and experimental virological assays to explore the molecular determinants behind the first detected HKU5r-CoV transmission to farmed mink^25^. We identify the loop2 region of the RBD as a key feature modulating receptor specificity and, in turn, host tropism of HKU5r-CoVs through direct interactions with the ACE2 receptor. A mutation at site 548 within nvHKU5r-CoV spike loop2 seems to be an adaptation to the virus’s new mink host. We show that substituting this same site with a naturally occurring polymorphism can enable the nvHKU5r-CoV RBD to utilize human ACE2. Together, our data highlight the role of the loop2 in shaping the zoonotic potential of these viruses.

Phylogenetic analysis places nvHKU5r-CoV within the HKU5-CoV-1 clade, clustering with bat derived viruses sampled from Zhejiang province, including WZ2 (Fig. 1a). When focusing on the spike and RBD regions, WZ2 is consistently the single closest known bat virus to nvHKU5r-CoV (Fig. 1b-c). This relationship is further supported by molecular dating of each genomic NRR, which reveal that WZ2 is the closest inferred ancestor in 16 out of the 47 NRRs (Table S4). These include parts of the spike (NRR30, NRR32, NRR33, NRR35, NRR37, NRR39), ORF3b (NRR41), ORF3c (NRR43), E and M genes (NRR45), and the N gene (NRR46), with tMRCA estimates ranging from 1903 to 2008 (Fig. 6b). These findings position WZ2 as the overall closest bat virus to nvHKU5r-CoV across the genome and suggest that, despite some of the tMRCAs being much older due to subsequent recombination events and unsampled virus diversity, the virus circulated in bats shortly prior to its detection in mink.

Recurrent recombination events define the evolution of nvHKU5r-CoV and its close bat relatives. Our analyses reveal that, within the RBD-encoding region of the spike gene, the loop2 region shows a distinct phylogenetic signal, where Q265 is the single closest relative to nvHKU5r-CoV (Fig. 2b-c). The relevance of Q265 to the evolution and emergence of nvHKU5r-CoV has previously gone unnoticed. Here, we show that this closest known loop2 virus can enable entry via nvACE2 similar to NV-SD-L1 and unlike any other known long loop2 HKU5r-CoV (Fig. 3a). Still, molecular dating of the loop2-encoding region (spike-NRR7) (Fig. 6c, Table S4) has relatively old tMRCA, with a median date in 1987 (95% HPD: 1950-2010). This means that the ability for these long loop2 viruses to efficiently use nvACE2 evolved many decades ago, and there is a wide unsampled diversity of closely related HKU5r-CoVs that can readily infect mink likely circulating in *P. abramus* bats. The potential of long loop2 viruses to use nvACE2 is further demonstrated by our full-length recombinant virus assay where HKU5-5 shows weak but detectable replication in nvACE2-expressing cells (Fig. 4d). One clear difference between the Q265 and nvHKU5r-CoV loop2 residues is a change from serine to arginine at site 548 (RQQLS in nvHKU5r-CoV and SQQLS in Q265; Fig. 2a,d). This seems to be an adaptive mutation selected for through circulation in mink, enabling the nvHKU5r-CoV RBD to directly engage the tyrosine at nvACE2 site 34, unique to this ortholog of the receptor (Fig. 5g).

The key role of the loop2 in modulating receptor engagement promotes large genetic diversity in this region, achieved through mutation, deletions, but also recombination of the loop2 between co-circulating HKU5r-CoVs. The loop2-specific phylogeny (Fig. 2c) reveals a single deletion event leading the short loop genotype. However, following this deletion, this shorter loop2 has recombined into multiple divergent HKU5r-CoV RBDs (Fig. 2c). This pattern may imply a selective advantage to this short genotype, although it is difficult to hypothesize what the underlying phenotype might be in the natural reservoir of these viruses, since both long and short loop2 RBDs exhibit comparable pseudovirus infectivity to the PabrACE2 (Fig. 3a). Regardless, these evolutionary patterns suggest that loop2 in HKU5r-CoVs is not solely defined by structural configuration or sequence similarity but is also shaped by broader evolutionary processes such as recombination, which may influence receptor compatibility and viral host range.

To dissect the molecular basis of different loop2 infectivity phenotypes, we performed mutagenesis experiments targeting notable loop2 residues (Fig. 5a,b). These results further imply that ACE2 usage is not dictated by loop2 length alone. For instance, HKU5-20S, a long loop2 virus, failed to use nvACE2 (Fig. 3a, Fig. 5a,b), but substitutions such as R543S and E550Q (Fig. 5b) modestly improved entry, suggesting that electrostatic tuning of the loop2 surface can enhance compatibility. Conversely, deletions mimicking short loop2 genotypes disrupted entry, indicating that truncation may compromise structural integrity rather than confer compatibility (Fig. 5b). These findings demonstrate that loop2-dependent phenotypes are shaped by distinct genotypic combinations, and that recombination or mutation can modulate these traits to potentially alter host tropism. These findings also suggest intermediate hosts like mink may play a significant role in honing human receptor compatibility for otherwise species-restricted viruses.

Among the mutations tested, R548S in the loop2 region of nvHKU5r-CoV stood out as a key determinant of human ACE2 usage (Fig. 5b). Interestingly, this mutation reverts the putative mink-specific adaptation of the nvHKU5r-CoV RBD back to the serine found in the majority of bat HKU5r-CoVs. Yet, the bat virus RBDs are unable to infect hACE2-expressing cells, including Q265 which shares the closest loop2 with nvHKU5r-CoV (Fig. 3a). This means that other residues within the loop2 or other parts of the RBD contribute to nvHKU5r-CoV’s unique ability to engage hACE2 when it has a serine at site 548. The molecular basis of this mutation’s effect on receptor usage lies on a direct interaction between spike site 548 and ACE2 site 34, the latter of which differs between humans and mink (Fig 5f,g). Importantly, this effect is observed without protease treatment (Fig. 5b), indicating that a single mutation, resulting from one change in the wobble-base of this codon, can directly enhance receptor engagement without relying on host proteases. The ability of this one naturally occurring mutation to unlock human ACE2 usage highlights the high zoonotic potential of HKU5r-CoVs. While currently unreported outside of bat and mink hosts, these viruses circulating in mink may be only a few genetic steps away from acquiring infectivity to humans or other intermediate hosts. Given the high-density and multispecies nature of fur farms^22,25^, such mutations could arise and be selected under the right conditions.

Interpreting our experimental results was largely aided by how well AlphaFold3 predictions of HKU5r-CoV RBD – ACE2 co-structures corresponded to the protein pairs’ infectivity phenotypes (Fig. 3, Extended Data Figs. 3-6). This was the case even when a single amino acid substitution in the input sequences was responsible for the phenotypic change, a prime example of this being the R548S mutation enabling nvHKU5r-CoV to use hACE2 (and correctly engage hACE2 in the AF3-predicted structure; Extended Data Fig. 9). Still, it should be noted that alignment between structural prediction and phenotype was not always the case, especially with RBDs more distant to the main HKU5-CoV-1 clade (Fig. 3g). Hence, current technologies for high-throughput structural predictions seem promising for assessing merbecovirus receptor specificity, as long as they are paired with experimental validation of the predicted interactions.

In summary, our findings paint a complex picture describing the proximal evolutionary history of the first HKU5r-CoV detected outside its natural bat reservoir. We highlight the spike loop2 as a crucial, but previously underappreciated, determinant of ACE2 engagement and host tropism. The phenotypic effect of mutations within loop2, however, largely differ between RBDs (Fig. 5a,b), meriting future research on the epistatic interactions between loop2 residues or even other parts of the spike. The bat virus genome that is overall most closely related (WZ2) and the genome with the closest loop2 to nvHKU5r-CoV were sampled in Zhejiang and Guangdong, respectively, two of China’s provinces where primary fur processing and wholesale facilities are located (Fig. 6d)^19^. It is unclear exactly how fur processing facilities are connected to the mink farms located in northern provinces, such as Shandong where nvHKU5r-CoV was sampled. Furthermore, the host range of *P. abramus* bats, which carry these viruses, extends across all these provinces. Hence, virus movement through farmed animal trading routes or transmission to mink from bats in Shandong are both possible routes for how nvHKU5r-CoV emerged in the mink. These findings highlight the importance of proactive surveillance at the interface between bat populations and farmed animals in order to prevent future zoonotic spillover events from this group of coronaviruses.

## Methods

### Biosafety

Prior approval for experiments using full-length recombinant HKU5 coronaviruses was obtained from the Institutional Biosafety Committee (IBC) of the University of North Carolina at Chapel Hill. All experiments using full-length recombinant HKU5 viruses were performed under Biosafety Level 3 conditions with personnel wearing full-body personal protective equipment and HEPA-filtered respiratory protection.

### Sequence retrieval and alignment

Full-length genome and Spike gene sequences were retrieved using BLASTn (NCBI; https://blast.ncbi.nlm.nih.gov/Blast.cgi), with the mink-derived HKU5r-CoV isolate (GenBase: C_AA075371.1) used as the query. To ensure inclusion of recently classified lineages, additional sequences were obtained from GenBase (https://ngdc.cncb.ac.cn/genbase/)^35^, including all available mink-HKU5 and HKU5-CoV-2 genomes^6^. Sequence acquisition was finalized on the following dates: HKU5-CoV-1 and related merbecoviruses from GenBank on October 24, 2024; mink-derived HKU5r-CoVs^18^ on October 24, 2024 (3 sequences); HKU5-CoV-2^6^ on March 14, 2025 (6 sequences); and MRCoV^25^ on May 21, 2025 (1 sequence) (Table S1). Initial BLAST retrieval yielded 100 full-genome sequences, which were curated to a final set of 44 genomes based on alignment quality, completeness, and removal of redundant or low-quality entries. For spike gene analysis, 88 sequences were retrieved and curated to 61 entries, with 10 additional sequences manually curated and appended (nvHKU5r-CoV and HKU5-CoV-2). Sequences were excluded based on quality control criteria including excessive ambiguous bases (>1% ‘N’ characters), poor alignment quality, and incomplete coding regions. The sequences were codon-aligned using PAL2NAL^36^, then one sequence (MT648449.1) was excluded due to alignment errors.

All nucleotide and amino acid alignments were performed using MAFFT v7.526^37^ with default parameters. For receptor-binding domain (RBD) analyses, the RBD region was extracted based on the annotated coordinates of the spike protein from *Pipistrellus bat coronavirus HKU5* (NCBI: YP_001039962.1) and aligned using codon-aware alignments derived from the spike gene dataset to preserve reading frame integrity for downstream phylogenetic and mutational analyses.

### Phylogenetic analysis

Maximum likelihood phylogenies were constructed using IQ-TREE v2.4.0^38^. Separate phylogenies were generated for full-length genome sequences, spike gene sequences, and RBD regions, analyzed at both nucleotide and amino acid levels. Model selection was performed using ModelFinder^39^ within IQ-TREE. The best-fit models were TIM2+F+R4 for nucleotide-based RBD alignments and JTTDCMut+R3 for amino acid-based RBD alignments, selected from a comprehensive model set including Blosum62, Dayhoff, DCMut, JTT, LG, Poisson, PMB, WAG, and EX variants. Branch support was assessed using 1000 ultrafast bootstrap replicates^40^. All trees were rooted using HKU5-CoV-2 as the outgroup and visualized using FigTree v1.4.4 (https://github.com/rambaut/figtree).

To account for the confounding effects of recombination, a recombination-free phylogeny was constructed. Recombination breakpoints were identified using Recombination Detection Program (RDP) v5.44 (RDP5)^24^ with the following parameters: linear sequence configuration, Bonferroni correction (p ≤ 0.05), original RDP5/4/3 decision tree, requirement for topological evidence, breakpoint polishing, and alignment consistency checks. Detection methods included RDP^41^, GENECONV^42^, Chimaera^43^, MaxChi^44^, BootScan^45^, SiScan^46^, and 3Seq^47^. Only recombination events supported by at least six independent methods out of seven were retained^3^. Recombinant regions were masked, and the resulting alignment was used to reconstruct the recombination-free ML tree (Extended Data Fig. 1c).

### Mutation mapping

Amino acid differences within the RBD of HKU5r-CoVs was visualized using Snipit v1.6^48^ with NV-SD-L1 designated as the reference sequence and compare it with its closest bat relative, WZ2, and the HKU5r-CoV reference genome, HKU5-1 LMH03f. The aligned RBD amino acid sequences were used to generate a residue-level comparative map, showing key substitutions across representative HKU5r-CoVs.

### Similarity plot analysis

Similarity plot analyses were performed using SimPlot++^49^. For whole-genome comparisons, NV-SD-L1 was used as the reference genome. The analysis was conducted using a 250 bp sliding window and a 20 bp step size, applying the Jukes-Cantor substitution model. For spike gene comparisons, the same parameters were used to calculate percent identity between NV-SD-L1 and representative HKU5r-CoVs. These plots were used to identify regions of high and divergent similarity, including those suggestive of recombination.

### Tanglegram

ML phylogenies were constructed for both the full RBD and the loop2 subregion using IQ-TREE v2.4.0^38^. The best-fit model for the RBD tree was TIM2+F+R4, and for the loop2 tree, TIM2e+R2, as determined by ModelFinder^39^. A tanglegram was generated using the phytools v2.4-4^50^ package in R v4.3.3 to compare the topologies of the two trees and assess phylogenetic incongruence between the RBD and loop2 regions.

### Recombination analysis

To investigate recombination across the HKU5r-CoV genome and Spike gene, we applied the Genetic Algorithm for Recombination Detection (GARD)^26,51^ implemented in the HyPhy software suite^52^. Prior to the analysis, redundancy was reduced using CD-HIT v4.8.1^53,54^ with a 99% identity threshold, resulting in 45 unique clusters for the whole genome and 55 for the Spike gene. Clustered sequences were re-aligned using MAFFT v7.526^37^.

GARD analyses were executed on a high-performance computing server using the MPI-enabled version of HyPhy^52,55^ (HYPHYMPI), with 20 parallel processes and the GTR substitution model. The environment variable TOLERATE_NUMERICAL_ERRORS=1 was enabled to ensure computational stability. Resulting breakpoint-partitioned alignments were used to define NRRs, which were subsequently used for ML phylogenetic reconstruction.

To extract individual NRRs from the full genome and spike-based alignment, we developed a custom python script that segments the alignment based on GARD-identified breakpoint coordinates. The script uses Biopython v.1.84^56^ to generate region-specific FASTA files and automatically invokes IQ-TREE v2.4.0^38^ to infer phylogenies for each region. For regions 1 through 13, trees were constructed using the GTR+F+I+R4 model with 10000 ultrafast bootstrap replicates. The final region, extending from the last breakpoint to the end of the alignment, was processed using the GTR+F+I+G4 model. All trees were midpoint-rooted and used for downstream analyses.

To identify the closest bat relatives of mink-derived HKU5r-CoVs across NRRs, we assessed monophyly and extracted sister taxa using the ete3 python package^57^. For each NRR tree, the script assessed whether the two mink sequences (C_AA075371.1 and C_AA075373.1) formed a monophyletic clade. If monophyly was confirmed, the most recent common ancestor (MRCA) node was used to extract sister taxa and calculate genetic distances.

Closest relatives were defined based on a genetic distance threshold of <0.05 and bootstrap support ≥80. To minimize redundancy, representatives were selected based on uniqueness and minimal co-clustering, ensuring that the closest relatives were phylogenetically and geographically informative.

### Cells and viruses

293T cells (ATCC CRL-3216) and BHK-21 (ATCC CCL-10) were maintained in DMEM (Gibco) supplemented with 10% FBS (GE Hyclone), penicillin-streptomycin (Gibco), and L-glutamine (Gibco) and maintained at 37□°C and 5% CO2. Cell line species origins were confirmed by cytochrome sequencing, and all cell lines were verified as mycoplasma negative with MycoSniff PCR Kit (MP Bio). Stable cell lines expressing ACE2 orthologs were produced as previously described^2,58^. Briefly, BHK-21 cells were infected with lentiviral transduction vectors expressing ACE2 orthologs and puromycin N-acetyl transferase. They were subsequently selected for using 1 µg/mL puromycin for approximately one week post-transduction. BHK cells were maintained under puromycin selection for the duration of the study. Alternatively, ACE2 orthologs were cloned into the sleeping beauty transposon plasmid, pSB-RFP-Puro (Addgene, #60513) and transfected along with transposase, pCMV(CAT)T7-SB100 (Addgene, #34879), into Vero81 cells lacking endogenous ACE2. Cells were stably selected with puromycin (10ug/ml) for 2 weeks.

The molecular clone derived HKU5-nLuc reporter viruses were generated as previously described^59^. Briefly, genomic cDNA sequences were ligated, in vitro transcribed, and electroporated into BHK cells. Stocks of recombinant HKU5-nLuc were then propagated in Vero81 cells stably expressing ACE2 orthologs at 37C with 5% CO2. Viral supernatants were harvested after extensive cytopathic effects were observed and clarified by centrifugation and stored at -80°C until use.

### Pseudotype production

Pseudotypes were generated as previously described^2,60–62^. 293T cells were seeded in poly-lysine treated 6-well plates and incubated at 37°C for 24 hours. Cells were then transfected with spike plasmids using polyethyleneimine (Poly Sciences. 24-hours post transfection, cells were infected with VSV-ΔG-pseudotype VSV particles at a multiplicity of infection of two in serum-free media and left to incubate for one hour. To remove seed particles, cells were then washed three times and left in serum free media. Pseudotypes were then collected at 48-hours post infection, clarified by centrifugation, aliquoted, and stored at -80°C until needed.

### Replication competent VSV

Replication competent VSV (rcVSV) NV-SD-L1 expressing GFP was generated as previously described^63^. Briefly, HEK293T cells stably expressing T7 RNA polymerase were transfected with plasmids encoding VSV N (Addgene #64087), P(Addgene #64088), L (Addgene #64085), G (Addgene #64084), and an antigenome copy of the viral genome (Addgene #31842) under control of the T7 promoter. Rescue supernatants were collected 24 hours-post transfection and clarified by centrifugation (5 minutes at 500 x g). Virus clones were plaque-purified and subsequently amplified on Vero81 cells stably expressing nvACE2 at 37°C with 5% CO2. Viral supernatants were harvested after extensive cytopathic effects were observed and clarified by centrifugation and stored at -80°C until use. Viral RNA was extracted from infected culture supernatants using the QIAamp Viral RNA Mini Kit following the manufacturer’s instructions. Reverse transcription and PCR amplification were performed using the SuperScript IV One-Step RT-PCR system, with primers flanking the exogenous glycoprotein insert (VSV.Forward: 5’-TAGTCTAGCTTCCAGCTTCTGA-3’ and VSV.Reverse: 5’-TCTCAAAATCGTGGACTTCCAT-3’). The resulting amplicons were gel-purified, and sequence verified using Oxford Nanopore sequencing (Plasmidsaurus Inc.). Sequencing data are available in the online supplementary repository (see “Data and code availability” section).

### Pseudotype entry assays

Entry assays were performed using standard approaches^2,64,65^. For mutagenesis experiments, 293T cells were seeded in black 96-well plates, transfected with receptor plasmids the next day, and infected 24 hours post transfection with equal volumes of viral pseudotypes. For the initial large HKU5 pseudotype panel, BHK-21 cells stably transduced with human ACE2, *Neogale vison* ACE2, or *Pipistrellus abramus* ACE2 were seeded in black 96-well plates and subsequently infected 24 hours later on ice with pseudotypes diluted with HBSS. Cells were washed once with cold PBS prior to infection with pseudotypes, centrifuged at 4°C, 1200 x g for 1 hour and incubated at 37°C overnight. Luciferase was measured using Bright-Glo reagent (Promega) approximately 18 hours post-infection. Relative entry was calculated by dividing raw luciferase values for each spike by the signal for no-spike pseudotypes. Plates were each measured 4 times to reduce error generated by plate reader background and analyzed individually. Relative entry values for all four replicates were averaged and plotted as heatmaps using Microsoft Excel.

### Plasmids

Plasmids used to generate chimeric spike pseudotypes, or *Merbecotypes*, were generated as previously described^2^. MERS-CoV/EMC12 spike (accession number: JX869059) was codon optimized for human cells, C-terminally FLAG tagged, and silent mutations were introduced near spike amino acids 341 and 617 to generate AflII and NotI restriction digest sites. Merbecovirus RBDs were also codon optimized, synthesized (IDT DNA), and used in downstream Infusion based cloning (Takara Bio) reactions to assemble chimeric spike proteins. Accession numbers for the spike panel can be found in Table S2. For receptors, *Pipistrellus abramus* ACE2 (GenBank GQ262782.1), *Homo sapiens* ACE2 (Q9BYF1.2), and *Neogale vison* ACE2 (GenBank XP_044091952.1) were synthesized (IDT DNA) and cloned into pcDNA3.1+ with NheI and ApaI sites.

### Full-length recombinant virus entry assay

Full-length recombinant HKU5r-CoVs encoding nLuc (HKU5-5 nLuc and NV-SD-L1 nLuc) were rescued by reverse genetics and propagated in Vero81 cells expressing PabrACE2 or nvACE2, respectively. For entry assays, Vero81 cells stably expressing ACE2 orthologs were seeded in 96-well black bottom plates at 2.5×10^4^ cells per well 24 hours prior to infection. Cells were inoculated with 800 PFU of nLuc viruses and subsequently incubated for 24 hours at 37 °C. At 24 hpi, cells were lysed and luminescence was quantified using the Nano-Glo Luciferase Assay System (Promega) according to the manufacturer’s instructions. Entry efficiency was calculated as RLUs normalized to the empty vector control within each experiment.

### Multi-step growth curves

Vero81 cells expressing nvACE2 or PabrACE2 were seeded in 6-well plats at 5×10^5 cells per well 24 hours prior to infection. Cells were infected with HKU5-5 or NV-SD-L1 at an MOI of 0.01 for 1 hour at 37C. After adsorption, cells were washed thrice with PBS and maintained in complete DMEM. Supernatants were collected at the indicated times post infection, and stored at -80C. Viral titers were determined via plaque assay on Vero81 cells expressing PabrACE2 as previously described^2^.

### Fluorescent focus forming assay

Vero81 cells stably expressing ACE2 orthologs were seeded at 2.5×10^4^ cells per well in black, clear-bottom 96-well plates 24 hours prior to infection. RcVSV stocks were 10-fold serially diluted in PBS in round-bottom 96-well plates, then transferred to the cell plates in duplicate and allowed to adsorb for 1 hour at 37C with 5% CO2. After adsorption, an overlay of OptiMEM supplemented with 1% carboxymethycellulose (CMC) was added, and plates were incubated for 24 hours. Cells were then fixed with 10% neutral-buffered formalin for 1 hour at room temperature, rinsed with distilled water, and air-dried. Fluorescent foci were imaged and quantified using a BioSpot reader.

### Electrostatic potential analysis

To determine the electrostatic surface potential for the RBDs and receptors, first the RBD amino acid sequences for HKU5-20s (381-614) or HKU5-21s (381-609) and the complete amino acid sequence for *Pipistrellus abramus* ACE2 were submitted to the AlphaFold3 online server^27^. The same was done for the NV-SD-L1 RBD (381-613) and the complete amino acid sequence for *Neogale vison* ACE2. Standard parameters were used for prediction. To analyze electrostatic potential differences, the APBS Electrostatics plugin^66^ was used in PyMol (Schrodinger Software: version 3.1.4) with standard parameters. For visualization, ACE2 amino acids after position 801 were trimmed to better compare with previously published structural data.

### Correlated sites analysis

To identify amino acid sites correlated with loop2 genotype (short vs. long), we first aligned the Spike protein sequences of HKU5r-CoVs using MAFFT v7.526^37^ and annotated loop2 genotypes based on the presence or absence of residues at alignment positions 550–553. Sequences with deletions at all four positions were classified as short loop2, while others were classified as long loop2. A codon-aware amino acid alignment was parsed into a site-wise dataframe using Biopython v.1.84^56^ and pandas v.2.2.2^67^. For each site, we performed a chi-squared test of independence between loop2 genotype. Bonferroni correction with an alpha of 0.05 was manually applied to adjust for multiple comparisons across all sites. Sites with corrected p-values below the significance threshold were considered statistically associated with loop2 genotype.

### Structural prediction modeling

Structural predictions of HKU5r-CoVs RBD in complex with ACE2 orthologs were performed using the AlphaFold3 server^27^. RBD sequences were trimmed based on conserved flanking regions identified in *Merbecovirus* Spike alignments, following the criteria described in our previous work^2^. Structural models with a predicted TM (pTM) and ipTM scores of ≥ 0.70 were accepted as reliable to reflect high-confidence structural predictions.

Predicted RBD–ACE2 co-structures were compared to experimentally determined complexes, including nvHKU5r–nvACE2 (PDB: 8ZWE)^25^, SARS-CoV-2–hACE2 (PDB: 6VW1)^31^, and PabrHKU5r–PabrACE2 (PDB: 9D32)^17^. All structures were visualized using UCSF ChimeraX^68^, and RBD–ACE2 interaction residues were identified using the contacts command. Residues within 4.0 Å of contact were defined as the interaction footprint.

ACE2 contact residues were extracted and processed using a custom python workflow. Structural parsing was performed using gemmi v.0.7.1^69^, sequence alignment with Biopython v.1.84^56^, and data handling with pandas v.2.2.2^67^ to map conserved binding sites across ACE2 orthologs. Benchmarking of AF3 predictions was performed using Boltz-1x (v.1.0.0)^29^, and chai-1 (v.0.6.1)^30^. The ‘predict’ module of Boltz-1x was run with default parameters, while the ‘fold’ module of chai-1 was used, utilizing templates and multiple sequence alignments retrieved from the associated server (‘--use-templates-server’, ‘--use-msa-server’). Both methods were run on four NVIDIA RTX 6000 GPU cards. All models were evaluated based on pTM and ipTM scores and visual inspection of interface resolution. AF3 models were used for all downstream analyses due to consistently higher confidence scores and more complete resolution of RBD–ACE2 interfaces.

### Molecular dating

Time-calibrated phylogenetic analyses were performed using BEAST X v10.5^70^. NRRs from the full-length genomes identified via GARD^26^ were used as input alignments. Sequence partitioning and XML configuration were conducted using the BEAUTi interface with the following parameters: site model set to GTR + Gamma equal weights), clock model set to uncorrelated relaxed clock (lognormal), and tree prior set to Coalescent Constant Size. Tip dates were parsed from sequence names using calendar date formats with variable precision. All other parameters were left at default settings.

Two independent MCMC chains were run per dataset, each with a chain length of 500,000,000 generations, logging every 100,000 generations. Priors were configured as follows: ucld.mean with a normal distribution (initial value = 1×10⁻³, mean = 1×10⁻³, standard deviation = 5×10⁻□), and constant.popSize with a lognormal distribution (μ = 6.0, σ = 0.5).

Convergence and effective sample sizes (ESS) were assessed using Tracer v.1.7.2^71^, with ESS values >200 considered acceptable. Final trees were summarized using TreeAnnotator^72^, with the target tree type set to HIPSTR^73^ and node heights set to common ancestor heights. Median node heights were used to estimate divergence times, including the time to the most recent common ancestor (tMRCA) between nvHKU5r-CoVs and their closest bat relatives.

### Temporal signal assessment

To assess whether NRR alignments had reliable temporal signal, we used Clockor2 v.1.9.1^74^ with the ML trees generated from GARD-partitioned full genome NRRs. A custom annotation file was created to define tip dates and lineage groupings for rate comparison.

### Geographic mapping

Sampling location information for representative HKU5r-CoVs was retrieved by surveying previously published literature describing virus isolation and characterization. Province-level geographic assignments were compiled and summarized in Table S1. To visualize the global distribution of *Pipistrellus abramus*, occurrence data were downloaded from the Global Biodiversity Information Facility (GBIF)^32^. Map visualization was performed using R v4.3.3, employing the rgeoboundaries v.1.3.1^75^, sf v.1.0-21^76^, ggplot2 v.3.5.2^77^, and ggspatial v.1.1.9^78^ packages. Administrative boundaries for China were retrieved at the first-level division (adm1) using geoboundaries(). The map was rendered using geom_sf() with scale and orientation annotations added via annotation_scale() and annotation_north_arrow().

### Statistics and data analysis

Each figure presenting cell entry data reflects results from four technical replicates. This number was selected to balance statistical robustness with experimental scalability. Experiments were conducted using different batches of pseudoviruses and cell preparations across multiple time points. Therefore, the data shown are representative of consistent trends observed across biological replicates.

## Data and code availability

Code and data used in this study are available on the following GitHub repository: https://github.com/TheSatoLab/nvHKU5_emergence.

## Acknowledgements

We would like to express our gratitude to all the members of Division of Systems Virology, The Institute of Medical Science, The University of Tokyo. We thank Dr. Jumpei Ito (The University of Tokyo) for providing insights on this work.

This study was supported in part by AMED ASPIRE program (25jf0126002 to Kei Sato); AMED SCARDA Japan Initiative for World-leading Vaccine Research and Development Centers “UTOPIA” (JP223fa627001 to Spyros Lytras, and Kei Sato); AMED SCARDA Program on R&D of new generation vaccine including new modality application (253fa727002 to Kei Sato); AMED Research Program on Emerging and Re-emerging Infectious Diseases (24fk0108907, 25fk0108690 to Kei Sato); AMED Japan Program for Infectious Diseases Research and Infrastructure (Collaborative Research via Overseas Research Centers) (25wm0225041 to Kei Sato); JSPS KAKENHI fund for the Promotion of Joint International Research (International Leading Research) (JP23K20041 to Kei Sato); JSPS KAKENHI grant-in-aid for Scientific Research A (JP24H00607 to Kei Sato); Mitsubishi UFJ Financial Group, Inc. Vaccine Development grant to Kei Sato; a grant from the Japanese Government Ministry of Education, Culture, Sports, Science and Technology Scholarship – Research Category (220235, to Jarel Elgin Tolentino).

Research results reported in this publication was also supported by the National Institute of Allergy and Infectious Diseases of the National Institutes of Health (NIAID/NIH) under Award Numbers R01 AI179720 to Michael Letko., R01 AI167966 and CEIRS Contract 75N93021C00014 to Ralph S. Baric., and T32AI007025 to Victoria Jefferson. The content is solely the responsibility of the authors and does not necessarily represent the official views of the National Institutes of Health.

## Author contributions

JET performed genomic, recombination, and phylogenetic analyses;

JET performed structural predictions;

JET, VAJ, and SL examined the structural predictions;

JET, VAJ, NJC, ML, and SL designed the experimental work;

VAJ performed pseudovirus assays;

NJC performed full-length recombinant virus assays;

JET and SL conceptualized the study;

RSB, ML, SL, and KS supervised the work;

JET, NJC, and SL wrote the original manuscript;

All authors reviewed and proofread the manuscript.

## Extended Data Figure legends

**Extended Data Figure 1. Expanded phylogenetic analysis of nvHKU5r-CoV and related merbecoviruses.**

**a.** Whole genome phylogeny including MERS-related CoVs (MERSr-CoVs), HKU4-related CoVs (HKU4r-CoVs), and both HKU5-CoV-1 and HKU5-CoV-2 lineages.

**b.** Spike gene nucleotide-based phylogeny showing consistent separation between HKU5-CoV-1 and HKU5-CoV-2, with nvHKU5r-CoV grouping within the HKU5-CoV-1 clade.

Both trees are rooted by the MERSr-CoV clade.

**c.** Phylogeny based on the RDP5 analysis recombination-free whole genome alignment of HKU5r-CoVs. The tree is rooted by the HKU5-CoV-2 clade.

Node support is annotated on all internal nodes in the trees. Scale bars represent nucleotide substitutions per site.

**d.** Bokeh-style heatmap visualization of genome-wide identity among HKU5r-CoVs.

**e.** Nucleotide percentage similarity between available nvHKU5r-CoV sequences for the whole genome (left), spike gene (middle), and RBD (right). Ambiguous nucleotides (‘N’) are not accounted in the calculations.

**Extended Data Figure 2. Recombination analysis of the Spike gene in NV-SD-L1 using GARD.**

**a.** Thirteen recombination breakpoints were identified across the Spike gene using GARD^26^. Breakpoint positions are mapped relative to the NV-SD-L1 genome.

**b.** ML phylogenies were reconstructed for each resulting non-recombinant region (NRR1–NRR13). Trees were rooted by the HKU5-CoV-2 clade. Node support is annotated on all internal nodes in the trees. Scale bars represent nucleotide substitutions per site.

**c.** Line graph showing the branch distance between nvHKU5r-CoV and its closest bat HKU5r-CoV in each spike-NRR tree.

**Extended Data Figure 3. Predicted spike–nvACE2 interaction footprints and experimental entry validation.**

AF3-predicted RBD–nvACE2 co-structures for all HKU5r-CoVs and control viruses. Structural footprints were compared to the native nvHKU5r–nvACE2 complex (PDB: 8ZWE)^25^. Viruses confirmed to enter nvACE2-expressing cells in pseudovirus assays are marked with a blue check mark; those that failed to enter are marked with a red cross. Entry phenotypes are based on experimental data from this study and previously published sources (see Table S2). Structural footprints were visualized using UCSF ChimeraX^63^, with residue contacts defined by a 4.0 Å cutoff distance.

**Extended Data Figure 4. Predicted spike–PabrACE2 interaction footprints and experimental entry validation.**

AF3-predicted RBD–PabrACE2 co-structures. Structural footprints were compared to the native PabrHKU5r–PabrACE2 complex (PDB: 9D32)^17^. Viruses confirmed to enter PabrACE2-expressing cells in pseudovirus assays are marked with a blue check mark; those that failed to enter are marked with a red cross. Entry phenotypes are based on experimental data from this study and previously published sources (see Table S2). Structural footprints were visualized using UCSF ChimeraX^63^, with residue contacts defined by a 4.0 Å cutoff distance.

**Extended Data Figure 5. Predicted spike–hACE2 interaction footprints and experimental entry validation.**

AF3-predicted RBD–hACE2 co-structures. SARS-CoV-2 bound to hACE2 is included as a positive control (PDB: 6VW1)^31^. Viruses confirmed to enter hACE2-expressing cells in pseudovirus assays are marked with a blue check mark; those that failed to enter are marked with a red cross. Entry phenotypes are based on experimental data from this study and previously published sources (see Table S2). Structural footprints were visualized using UCSF ChimeraX^63^, with residue contacts defined by a 4.0 Å cutoff distance.

**Extended Data Figure 6. Benchmarking of spike-ACE2 interaction models across prediction software.**

**a.** AF3 predictions of Spike–ACE2 complexes across HKU5r-CoVs and related merbecoviruses. Mean predicted TM-score (pTMmean) and interface predicted TM-score (ipTMmean), along with standard deviations are shown. Viral entry was defined by luciferase activity (Fig. 3a), with ≥10 units of relative entry indicating successful entry. **b-c.** Comparative benchmarking of AF3 against Boltz-1 (b) and Chai-1 (c) using identical input sequences.

**Extended Data Figure 7. Structural and sequence correlates of ACE2 usage in HKU5r-CoVs**

**a.** Amino acid residues at spike sites where short loop2 and long loop2 consensus sequences differ. Sites where the NV-SD-L1 matches the long loop2 consensus are colored in orange. Sites where the NV-SD-L1 matches the short loop2 consensus are colored in yellow.

**b.** Electrostatic surface potential of representative HKU5r-CoV RBDs in complex with ACE2 orthologs, visualized in PyMOL. Differences in charge distribution across loop2 highlight structural features associated with receptor engagement.

**Extended Data Fig. 8. Predicted RBD–ACE2 interaction footprints of HKU5-5 across species.**

AF3 structural models of HKU5-5 spike RBD in complex with ACE2 orthologs from *Neogale vison* (nvACE2), human (hACE2), *Pipistrellus abramus* (PabrACE2), and *Mus musculus*. The native RBD footprint from each respective PDB structure (PDB: 8ZWE for nvHKU5r-CoV–nvACE2; PDB: 6VW1 for SARS-CoV-2–hACE2; PDB: 9D32 for PabrHKU5r–PabrACE2; PDB: 7UFL for SARS-CoV-2-*Mus musculus* ACE2^79^) is shown in yellow on the ACE2 surface for reference.

**Extended Data Figure 9. Predicted hACE2 engagement by HKU5r-CoV spike mutants**

AF3 structural models of representative HKU5r-CoV RBD mutants tested against hACE2. The RBD footprint of SARS-CoV-2 (PDB: 6VW1) is shown in yellow on all hACE2 structures for reference.

## Supplementary Tables

**Table S1.** List of all HKU5r-CoV genomes used for phylogenetic analysis.

**Table S2.** List of ACE2-RBD Alphafold3 predictions with confidence scores (pTM, ipTM), predicted ACE2 contact residues (cutoff distance of 4.0Å) and entry phenotypes.

**Table S3.** List of interacting residues between the NV-SD-L1 R548S mutant RBD and hACE2 as inferred based on the AlphaFold3 prediction of the proteins’ co-structure (cutoff distance of 4.0Å).

**Table S4.** Summary of molecular dating estimates across the 47 whole genome NRRs and the 14 spike NRRs.

