## Extended Data Figures for "Structural and phenotypic plasticity of the RBD loop2 region is a key determinant for HKU5r-CoVs’ emergence in mink"

### Extended Data Figure 1

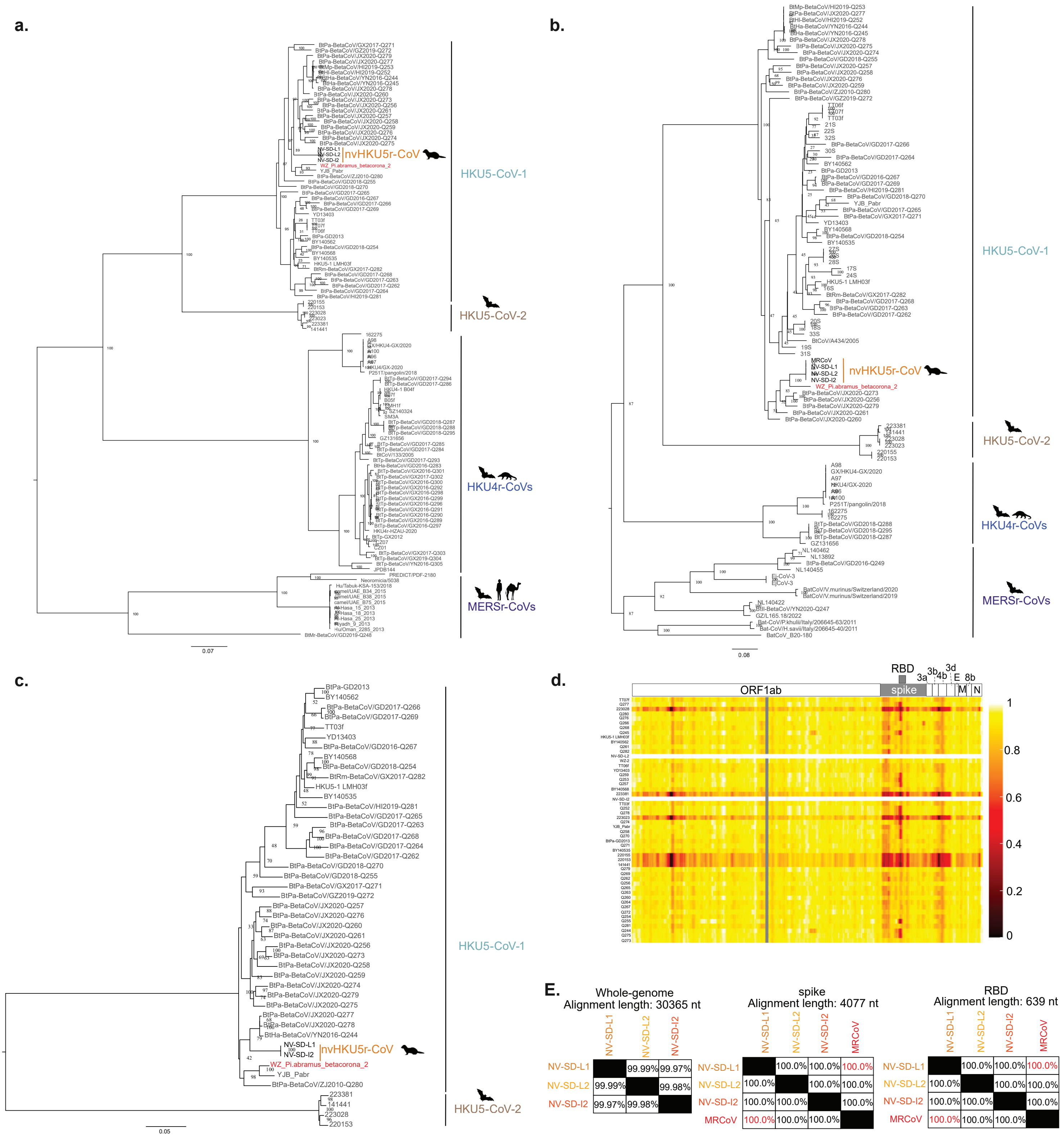

### Extended Data Figure 2

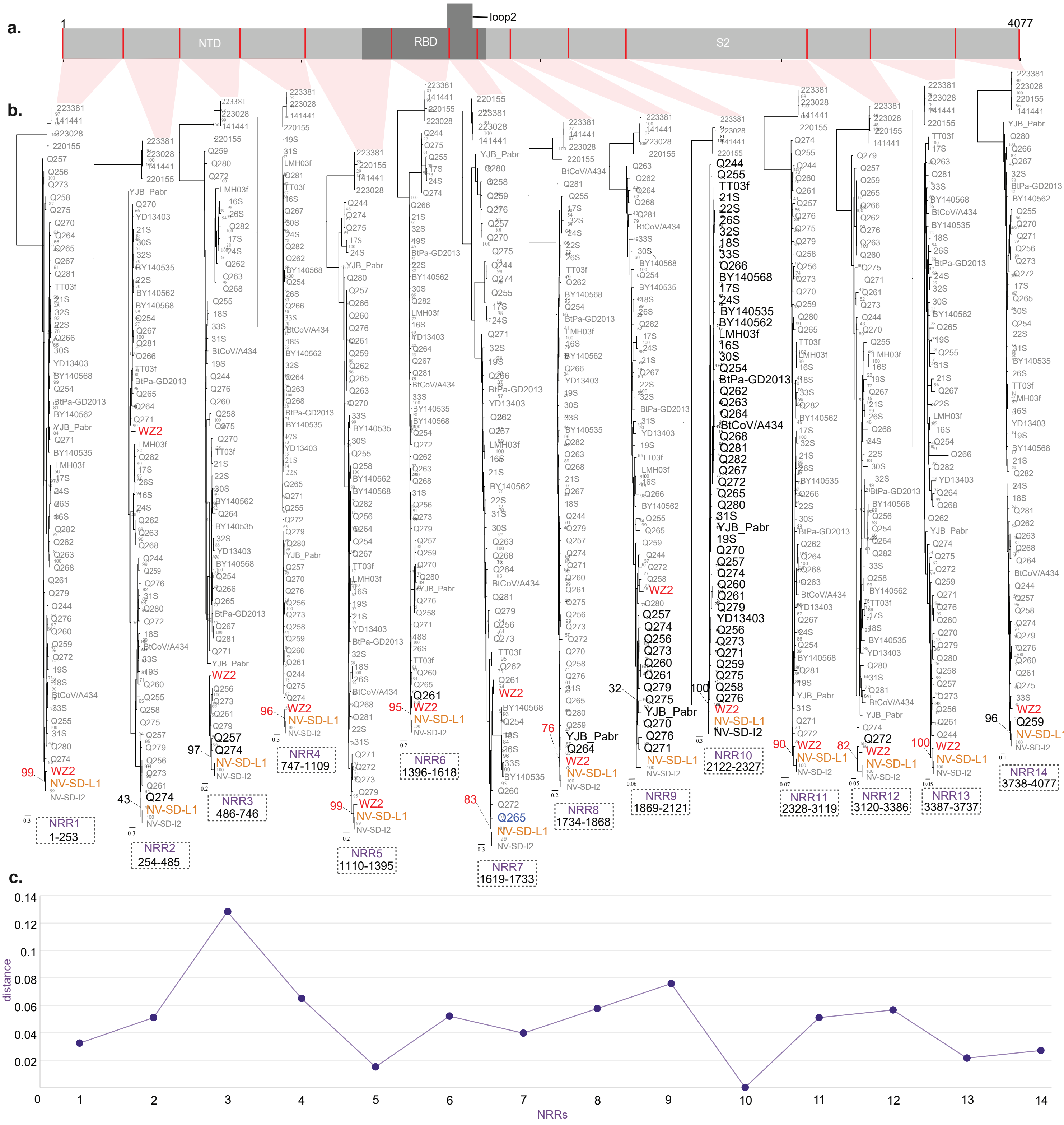

### Extended Data Figure 3

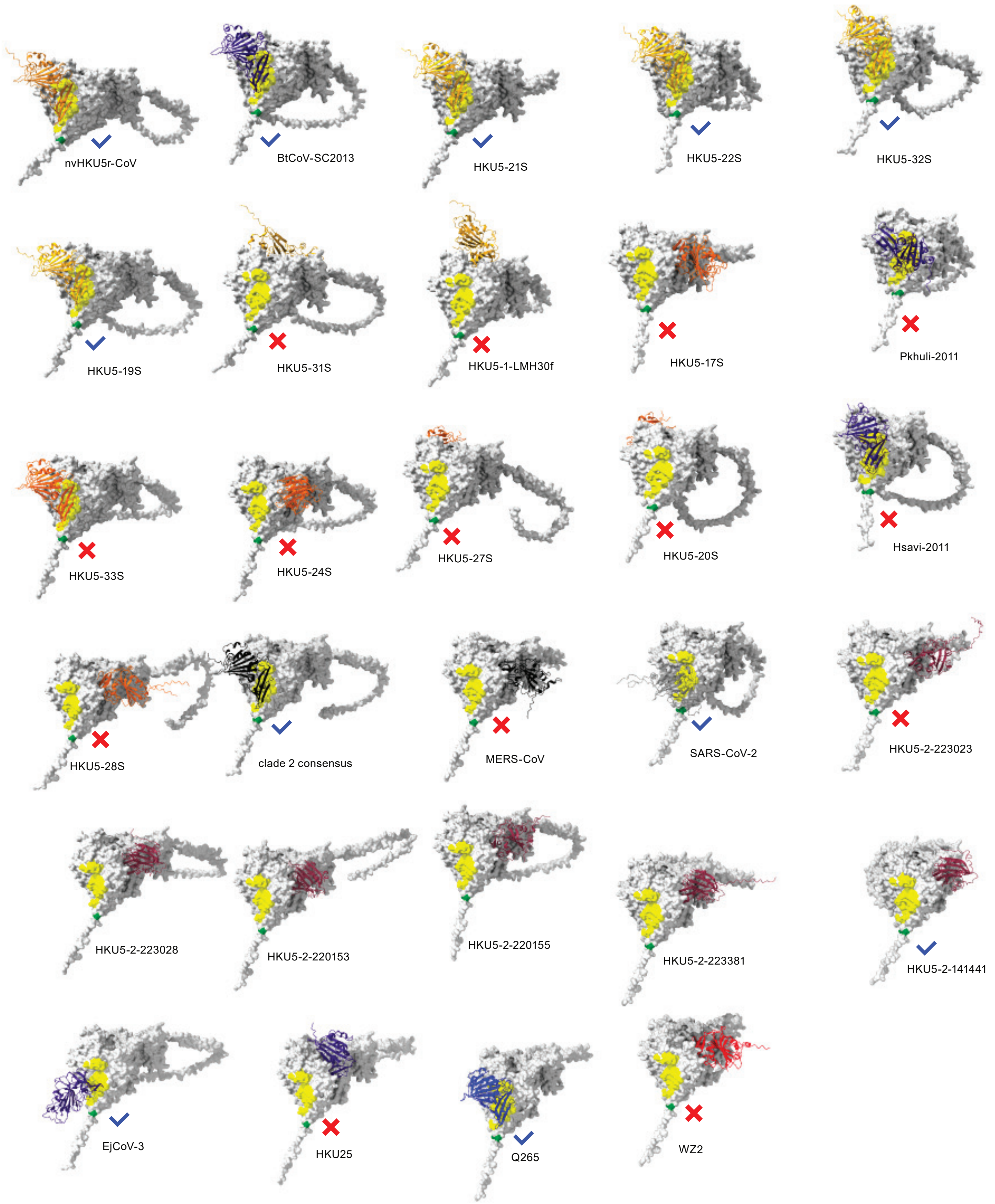

### Extended Data Figure 4

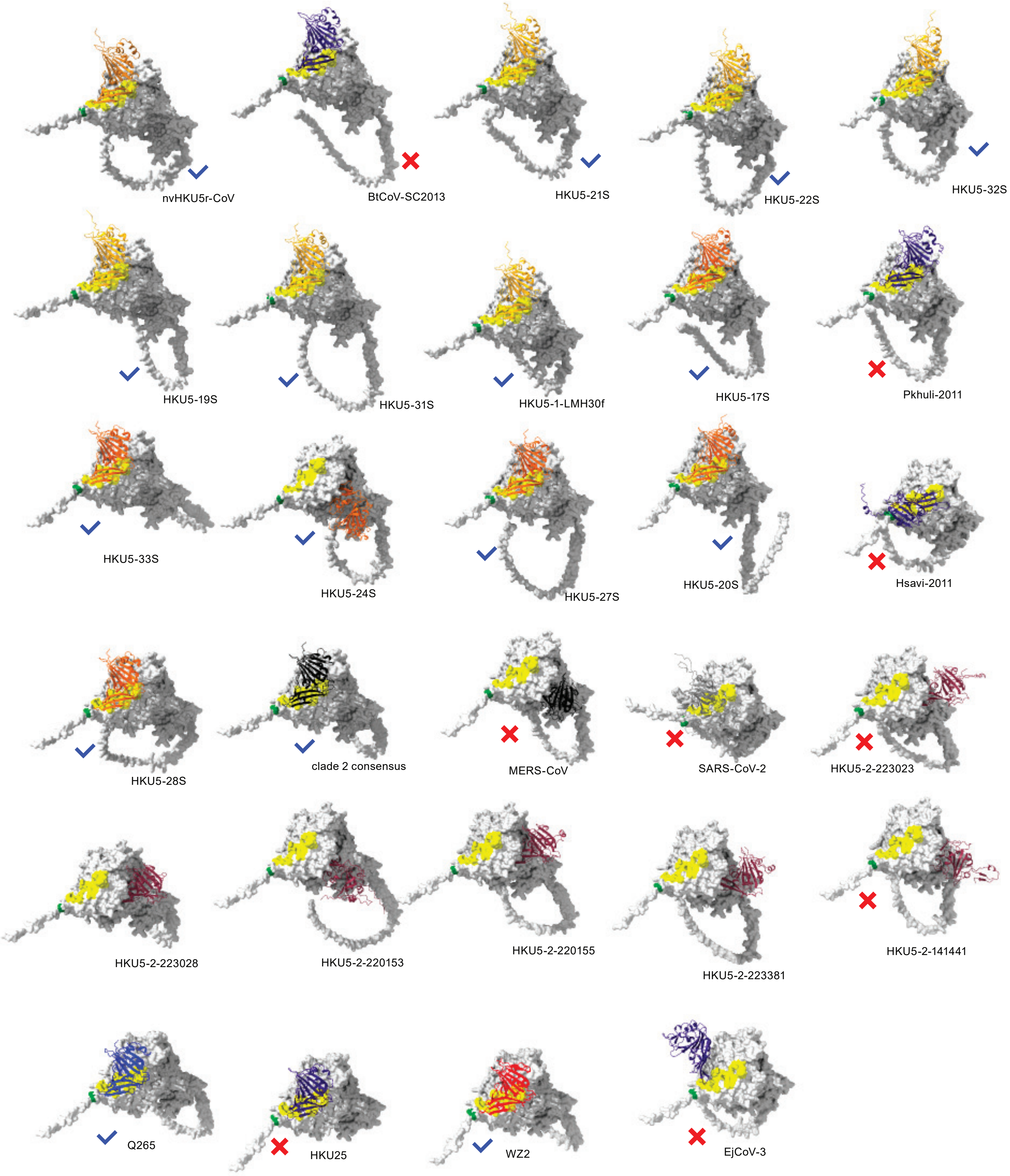

### Extended Data Figure 5

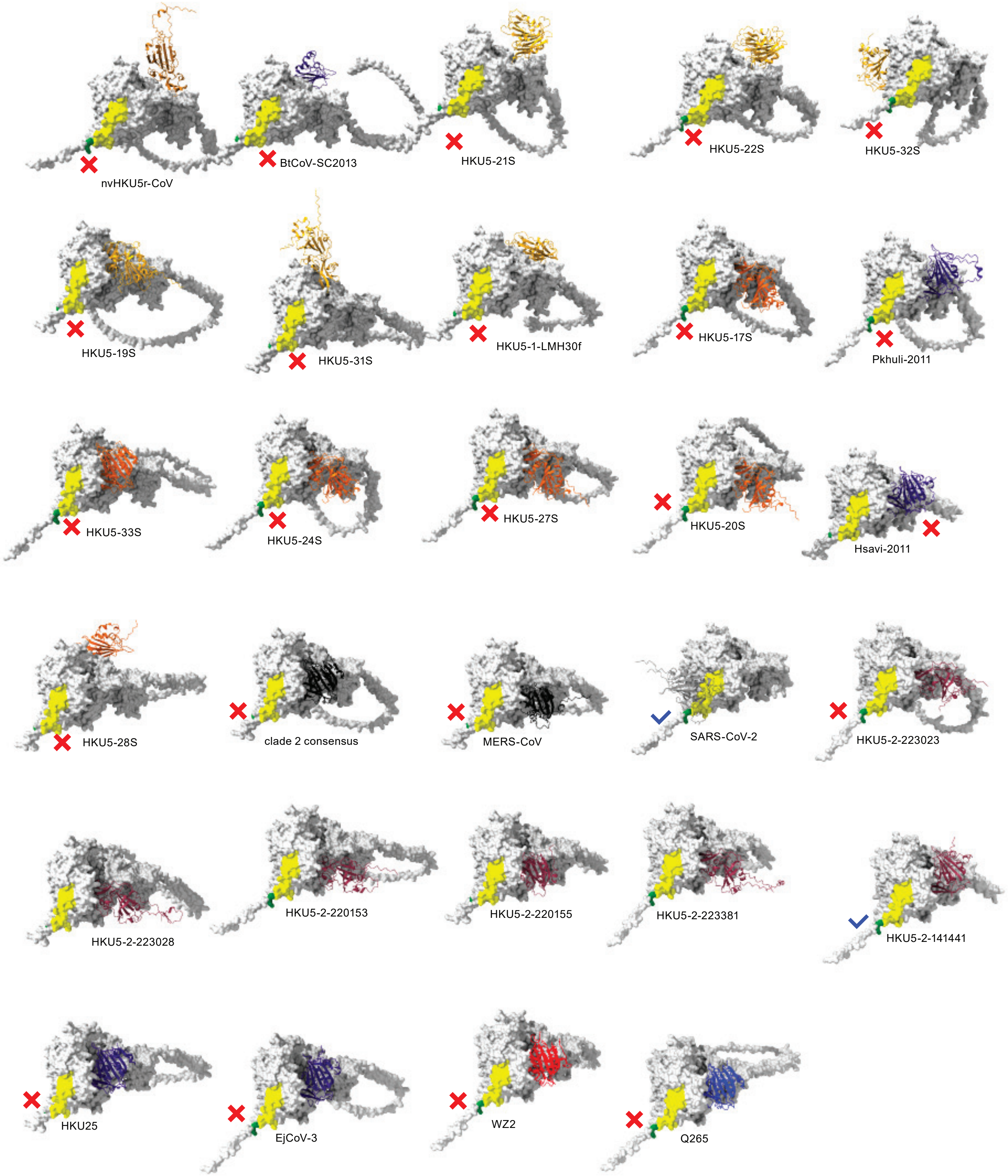

### Extended Data Figure 6

**a.** AlphaFold3

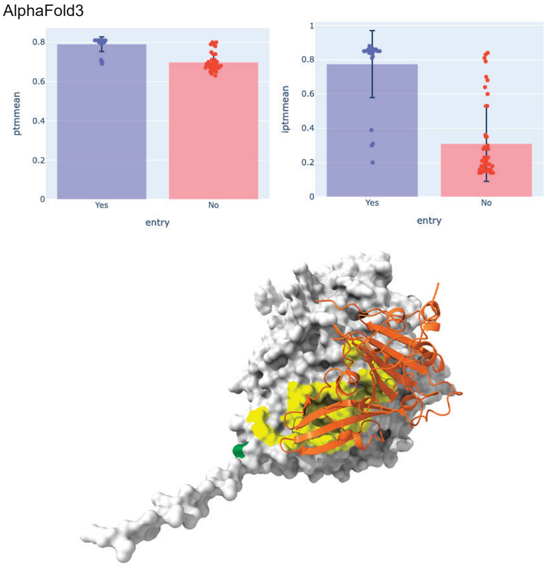


ptm= 0.80 iptm= 0.82

**b.** boltz1

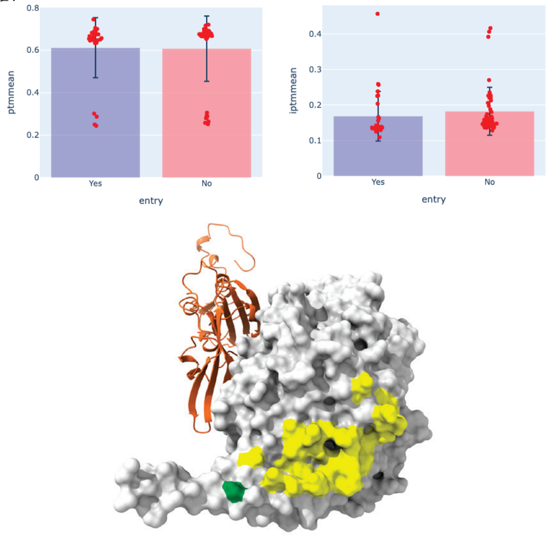


ptm= 0.68 iptm= 0.20

**c.** chai1

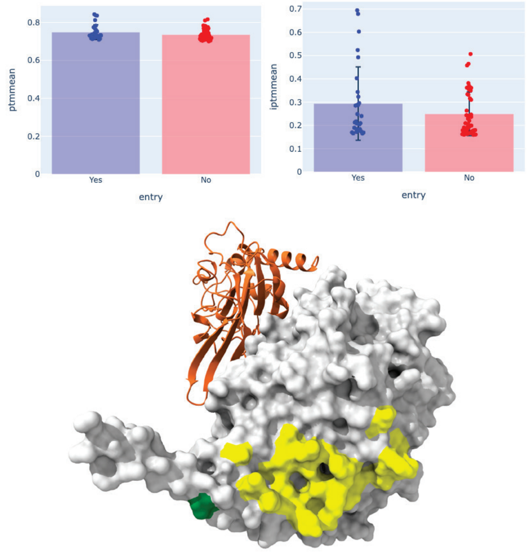


ptm= 0.73 iptm= 0.18

### Extended Data Figure 7

**a.** correlated sites

| alignment position | short-loop consensus | long-loop consensus | NV-SD-L1 sites |
|---|---|---|---|
| 506 | A | V | V |
| 507 | Y | H | H |
| 514 | T | S | G |
| 515 | S | T | T |
| 543 | T | R | S |
| 544 | K | D | D |
| 548 | Y | R | R |
| 549 | — | Q | Q |
| 550 | — | E | Q |
| 551 | — | L | L |
| 552 | — | P | S |
| 554 | — | F | — |
| 564 | I | V | V |
| 569 (NV-SD-L1 557) | G | S | G |

**b.** electrostatic potential

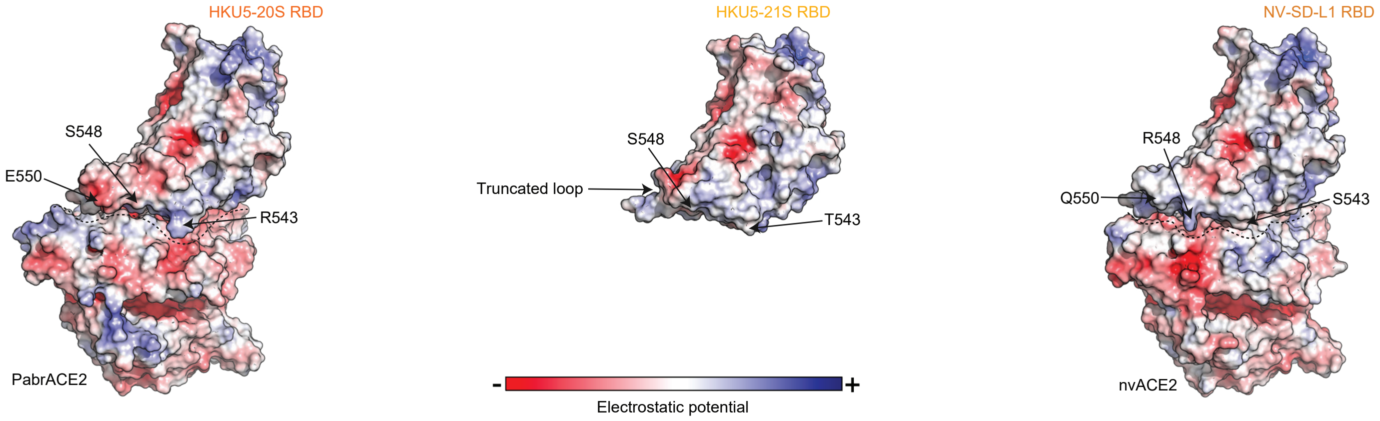

### Extended Data Figure 8

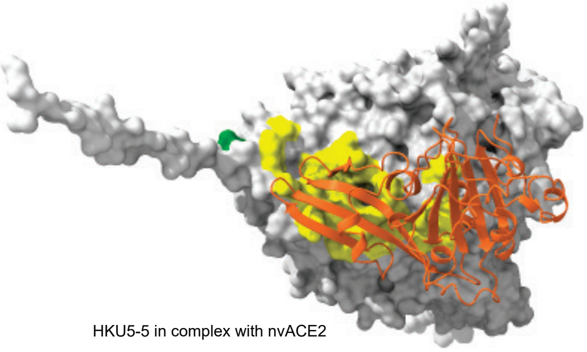

HKU5-5 in complex with nvACE2

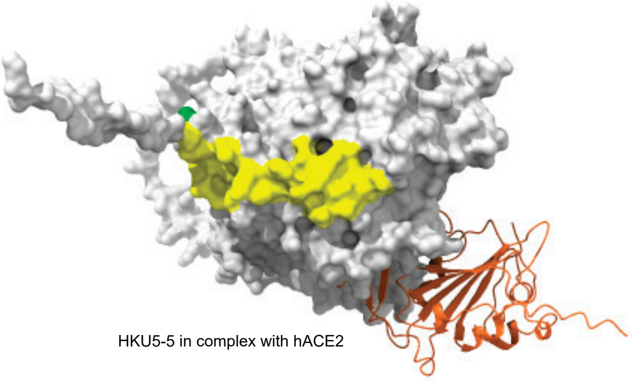

HKU5-5 in complex with hACE2

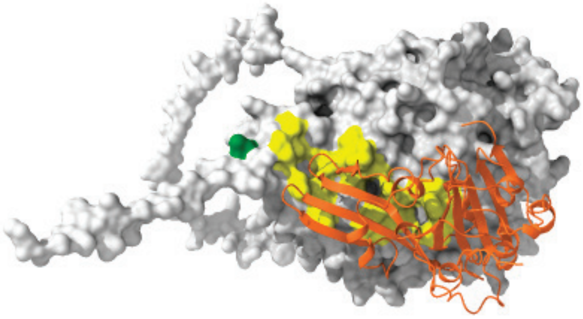

HKU5-5 in complex with PabrACE2

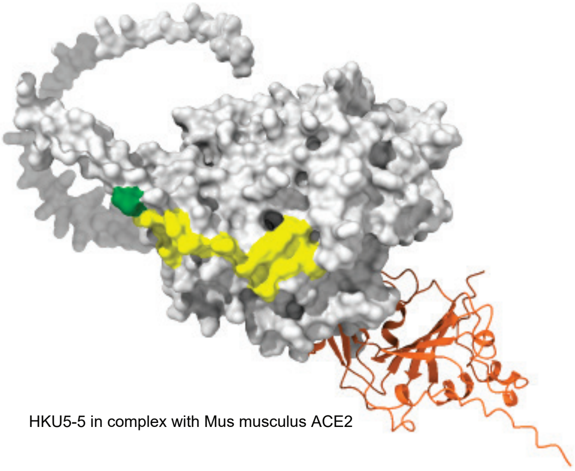

HKU5-5 in complex with Mus musculus ACE2

### Extended Data Figure 9

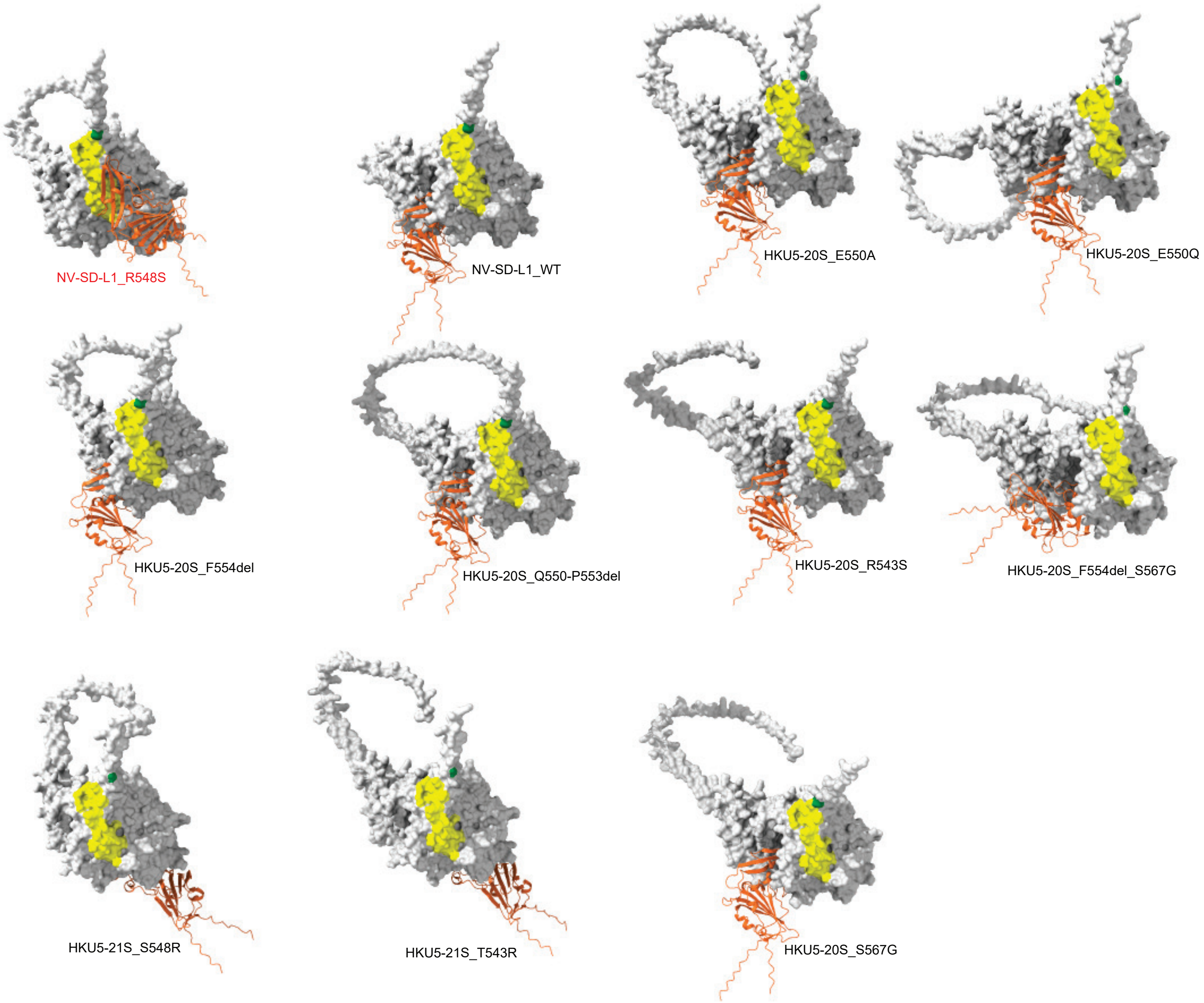
